# Low-latency multicamera 3D tracking of insects with Braid

**DOI:** 10.64898/2026.08.21.745392

**Authors:** Michael J.M. Harrap, Andrew D. Straw

## Abstract

Advances in camera technology and computer vision techniques have allowed researchers to track animals in 3D in ways which previously were difficult or impossible. Many such 3D tracking tools make use of multiple cameras, but unfamiliarity with the principles and technology involved can make it difficult to employ such techniques. In this protocol, we describe Braid, open-source software for live, multi-camera 3D tracking of insects. Using background-subtraction, Braid performs detection of objects without requiring the use of physical markers affixed to the insect. Braid constructs low-latency 3D position estimates using Kalman filtering and nearest neighbor data association. We document in detail the process of tracking freely flying bees within a flight arena using Braid. This protocol includes instructions on installation, configuration of cameras, setup, calibration, and operation. Within the system described here, we demonstrate that Braid can achieve position estimates accurate to <1 millimeter (within a 0.3 cubic meter volume). These factors make Braid suitable for tracking small, fast-flying animals like insects. Braid’s low latency allows live tracking, removing the necessity to collect large video files and making it suitable for integration in closed loop systems such as virtual reality. Code is available at https://github.com/strawlab/strand-braid

**Key features:** Guides the reader through setup and use of Braid multi-camera acquisition and tracking software.

Braid allows low latency, marker-free real-time 3D tracking of multiple objects, with a web-based GUI and integration with Rerun visualization software.

Braid has all necessary calibration tools incorporated within it. We provide detailed instructions on how to calibrate a Braid-multicamera system.

Braid is useful for 3D tracking of animals, robots, or other moving objects, particularly when these objects are small such as insects.

**This protocol is used in:** The components of the Braid system itself described here (cameras, PC, switch, etc.) was used as the ‘static camera’ system in Vo-Doan et al. [1]. Braid itself has also been used in: [2–8]. The algorithms used in Braid are similar or identical to those of its predecessor software Flydra [9] that has been used in: [10–19].

**Graphical overview:** **Graphical Overview:** Schematic of 3D tracking with Braid described here. Sequence of events that occur at each timepoint are described left to right. Red lines indicate the flow of information. Initially the flight of an insect, here a honey bee (inset), is observed by a series of synchronized cameras. The bee’s full flight trajectory to the yellow artificial flower is indicated by green points. Image frames from each camera are acquired and processed online with Strand Camera, with one instance of Strand Camera software running per physical camera. Central images show the image captured for the timepoint indicated by the black triangle on the bee’s flight trajectory. Strand Camera detects the insect in two dimensions. Green dots, visible within magnifications of image frames, indicate the location of these 2D detections. These detections are given to a single instance of the Braid software. Braid then utilizes a Kalman filter based algorithm and camera calibrations to produce estimates of the bee 3D position in real time. The full flight trajectory of the bee looking down the x (red axis), y (green axis), and z (blue axis) axes are shown (panels x, y and z respectively).

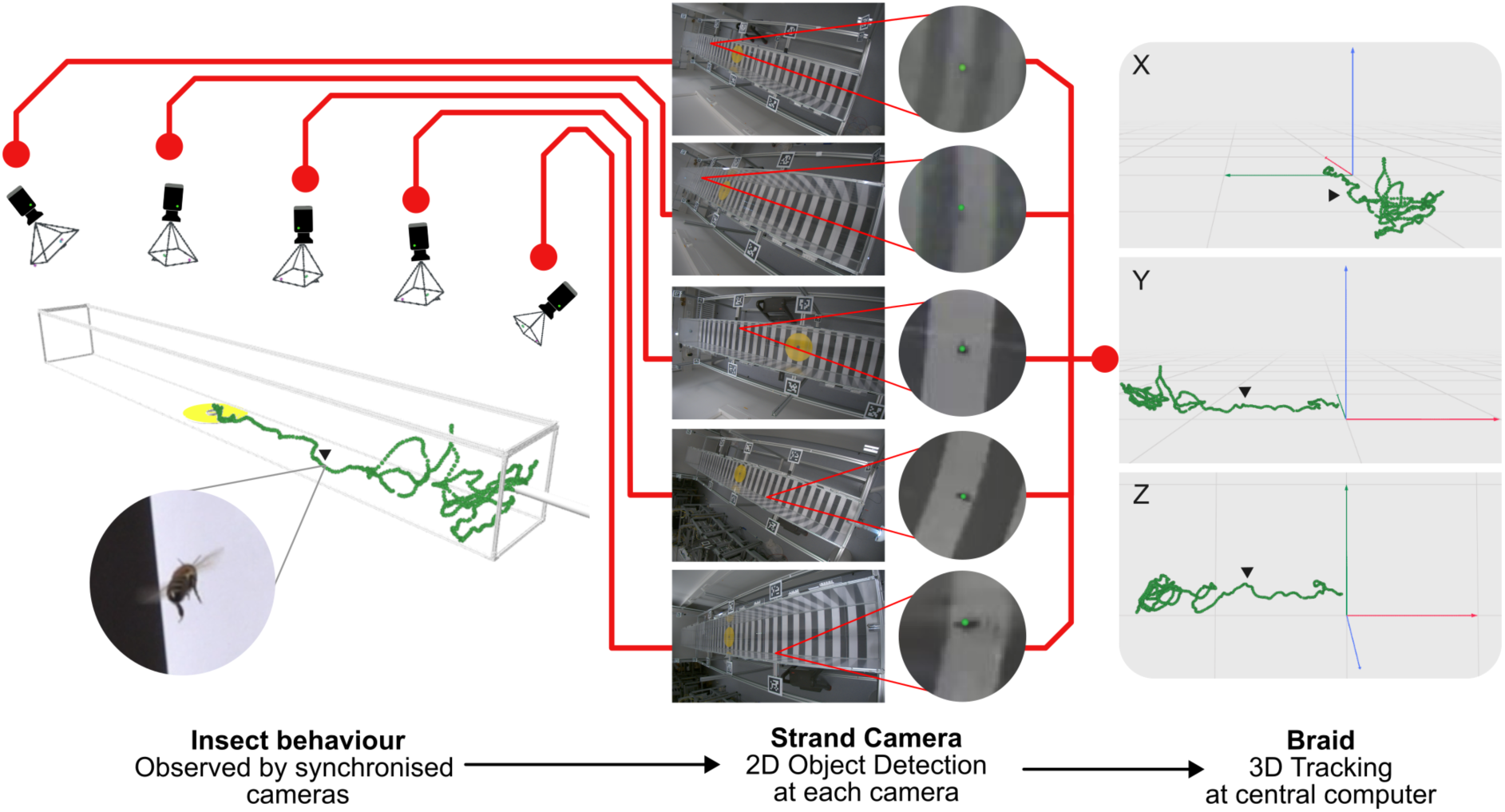

## Background

Many biological questions require quantification of animal subjects’ position and motion. Computer vision based tracking techniques allow researchers to automatically detect and track subjects (e.g. [13,20–26]). In recent years such computer vision tools have proliferated, especially with the advent of machine learning based techniques to facilitate object detection and classification (e.g. [27–29]). Automated tracking has been utilized to address a variety of biological questions. These include questions related to: flight behavior [30–33]; animal activity levels [34]; how animals behave in large groups [35–38], respond to stimuli associated with food [39–42], locate and pursue food [43–48]; avoid threats [49–51]; and navigate in complicated environments [52–61], among many others. Use of automated tracking tools can allow researchers to more quickly acquire large behavioral datasets than they would using manual identification [13] and can allow collection of quantitative data on behaviors which might have been represented in a more qualitative manner in the past allowing more detailed exploration and analysis (e.g. compare data collected in [62,63] with [64,65]).

How ‘position’ is represented in tracking applications can vary. The simplest applications track animals in 2D, often by cameras looking down a single axis, usually from above. Here, the animal’s movement is assumed to remain in a flat plane (e.g. [39,53,59–61]). This can be suitable for terrestrial animals where vertical position is comparable to ground level. However, tracking of animal position in three dimensions can give more complete insight, especially for flying animals. Yet flying animals may move quickly in three dimensions within relatively large areas, may be physically small, and sometimes can only be observed at a distance [26]. This is particularly true of insects. These factors are also true of swimming animals, which similarly move in 3D within a fluid environment [16]. While possible (see [66–71]), there are limitations to how well animals can be tracked in 3D using a single camera. If subject size varies, estimation of the animal’s distance from a single camera can be challenging. When animals are small and tracking volumes are comparably large, spatial resolution of cameras can become a problem. High temporal resolution cameras, needed to track fast-moving animals, often have more limited spatial resolution (i.e. to record images at a high frame rate, the cameras often have reduced resolution, i.e. smaller the image size). Additionally, occlusion can become an issue in some applications, particularly if the target organism is small. For these reasons, many 3D tracking systems make use of multiple cameras [1,9,13,17,22–24,26,72]. However, use of multiple cameras presents further challenges even when issues such as the calculation of 3D trajectories and data association across time are handled by existing software packages, such as setup, calibration and synchronization of cameras. Thus, without relevant familiarity with the steps involved, researchers can find implementation of multicamera tracking tools challenging.

In this protocol, we aim to address this difficulty while introducing the software Braid. Braid is free, open-source multi-camera acquisition and tracking software. A Braid system is made up of multiple synchronized instances of the single-camera based object detection software Strand Camera, which are coordinated by the Braid program itself. An instance of Strand Camera operates and controls an individual camera and performs “2D detection,” measuring the locations of the tracked object(s) in each camera’s field of view. Strand Camera’s object detection algorithm is based on background subtraction. A background model for the appearance of each pixel is made, and incoming images are compared with the model, allowing marker-less object detection. The 2D locations of objects found by each instance of Strand Camera are transmitted to the Braid program, which implements 3D tracking, creating estimates of the object’s position and velocity. Braid makes use of an extended Kalman filter and nearest neighbor data association algorithm [9]. This can be thought of as Braid modeling the 3D position and velocity of objects at each frame captured by the synchronized cameras. This model is then updated with each incoming frame based on the 2D detections from individual cameras (see [9] for detail on the Kalman Filtering applied in Braid). Together, the system is robust to brief occlusion of the object being tracked, individual camera detections can still contribute to the position estimates even when only one camera can detect the object, and uncertainty of the estimates is maintained. Braid’s object detection is robust, and not target specific, thus Braid can be applied to track any object in 3D space but is particularly useful in tracking small, fast-moving targets such as insects. Braid’s low latency allows live tracking of objects so 3D tracking data to be recorded without the need for video to be saved, vastly reducing the file size and storage requirements needed for 3D position data collection. Braid is a successor to Flydra [9], being at its core a Rust-language based rewrite of Flydra. Thus, the details of object detection, data association and application of the Kalman filter to track object’s position and velocity remain similar or identical to those described for Flydra in Straw et al. [9]. Like Flydra, Braid is capable of tracking multiple objects simultaneously and also tracking underwater objects from cameras above the water by modeling the refractive index boundary [16].

Here we describe the setup of the basic Braid system, where cameras are synchronized with precision time protocol (PTP) over a gigabit ethernet network. The protocol demonstrates use of Braid in the context of tracking bees within a tunnel flight arena (in a setup similar to that performed in [73–76]). Our goal is to cover all steps with detailed descriptions of what each step entails in a manner that can be understood and followed without extensive prior experience of computer vision or 3D tracking tools.

## Materials and reagents

### Biological materials

1. Honey bees, *Apis mellifera (L.)*. Four hives were maintained onsite using standard beekeeping practices, allowed free access to forage in the local environment. These bees were trained to visit the lab via a window-access (see equipment item 20) providing 1M sucrose (see below and Supplementary material 2).

### Solutions

1. 1M Sucrose, as a reward for artificial flowers.
2. 30% ethanol solution (by volume) for cleaning.

### Equipment

1. a PC with Linux Ubuntu 24.04 amd64 installed, the full specifications of the PC we use are provided in Supplementary material 1.
2. Work surface (Fetra Fechtel Transportgeräte GmbH, Borgholzhausen Germany, Item name: Rolltisch, 150kg, article number: 5864), to place the PC upon.
3. Room dividers (KAISER+KRAFT GmbH, Stuttgart Germany, Item name: Info-Wand Eurokraft basic, article number: 759462 49) two are required to obscure PC and experimenters from cameras during operation.
4. Ethernet cables, six required (S-Impuls Handels, Markt Bibart, Germany, model name: Patchcable, 7.5m, cat6A S/FTP, article number: 75717)
5. 10G Gigabit ethernet cable, one required (Ubiquiti, New York NY, USA, Model: 10G Base Direct Connect Cable - SFP+ to SFP+ - 3 m - 6 mm - passive)
6. Gigabit ethernet switch (Hewlett Packard Enterprise, Spring Houston, TX, USA. Model: HPE Networking Instant On GigE Switch Series 1930 24G Class4 PoE 4SFP/SFP+ 195W serial number: JL683B)
7. five Basler GigE cameras (Basler, Ahrensburg, Germany. Model: a2A1920-51gcBAS GigE camera with the Sony IMX392 CMOS sensor)
8. Lenses (Edmund optics, Barrington NJ, USA. Model: 4mm UC Series Fixed Focal Length Lens), five required
9. Ball mounts (PanaVise, Reno NV, USA. Model: 851-00 Knuckle Adjustable Knob and Screw) with T-nut fittings and screws to attach to the support and camera mounting frame (M5 T-nuts, Item Industrietechnik, Solingen, Germany, article number 0.0.480.54 and an M5 25mm long countersunk screw, ISO 10642). Nine are required in total (five are required for a camera mounting, a further four are required for light arrays).
10. Mounting plates to attach Basler cameras to the ball mounts. These have holes positioned to match the 3mm screw fittings of the cameras and have a socket for a ¼” nut. These custom parts are 3D printed (Prusa Research a.s., Prague, Czech Republic, Printer: Prusa i3 MK3S, Material: Prusament PETG Jet Black). CAD file is provided in [77], at: *protocol_downloads/custom_equipment_cad_files/3D_printed_parts/ace2-gige_mount.stl*.
11. Checkerboards made of black and white squares for intrinsic camera calibration. Two 9 by 7 square checkerboards are used, one with 30mm squares, one with 10mm squares. All checkerboards were produced via the Python script in the Strand-Braid repository https://github.com/strawlab/strand-braid/blob/main/docs/user-docs/scripts/draw_checkerboard_svg.py. Each is printed out full size, stuck flat (with no folds, bumps or creases) to a 5mm thick piece of foamboard. To the back of the foamboard a handle for the checkerboard made from a plastic cup or more foamboard (depending on the board) is stuck with tape
12. AprilTag plates. Made using ‘L204 White, black, white 1.6mm trolaze’ (Trotec Laser Gmbh, Marchtrenk, Austria. Catalog number: 91640 1031005903397) onto which the AprilTag designs are etched and cut free with a CO2 laser cutter (Trotec Laser Gmbh, Marchtrenk, Austria, model: Speedy 100). CAD files are provided in [77], at: *protocol_downloads/custom_equipment_cad_files/AprilTag_plates/*.
13. Sloped mounting blocks, to angle position of AprilTags, four required. These are 3D printed (using materials as above). These are adhered to AprilTag plates with the trolaze adhesive and to the Support and camera mounting frame with double sided tape. CAD files are provided in [77], at: *protocol_downloads/custom_equipment_cad_files/3D_printed_parts/tag_grove.stl*.
14. Cable ties, to attach camera cables to the flight arena support frame (OBI Group Sourcing GmbH, Wermelskirchen, Germany, Item name: Kabelbinder Weiß, article number: 3511110)
15. Light arrays. To illuminate the arena. Four simple light arrays are used. Each array is made of a pair of LED light units (LED Umrüst-Modul, Eltric K. Heckel GmbH, Bayreuth, Germany, article number: 541135) screwed to a steel plate. These are in turn attached to the support frame using a ball mount (see above). Light arrays are wired together in parallel and powered via mains power supply (wire: Huber+Suhner, Herisau, Switzerland, article: Radox 125, 1mm, RED and BLUE; connectors: Wago, Minden, Germany, article: Verbindungsklemme 221-413; plug socket and switch: Eltric K. Heckel GmbH, Bayreuth, Germany, Anschlussleitung mit Schukostecker und Zwischenschalter, article number: 026000). Circuit diagrams for each array and connection of lights to mains are available in figure S5 and S6
16. Screwdrivers (Wiha-Softfinish, Schonach, Germany, EAN code: 4010995010218) and hex key sets (Ifixit hex key set, San Luis Obispo CA, U.S.A, article number: IF145-859-2).
17. Insect flight arena, made from Makerbeam 10x10mm aluminum profile extrusions and associated fittings for this profile system (Makerbeam B. V., Utrecht, Netherlands) and UV permeable acrylic glass of 3mm thickness (Rohm and Haas, Philadelphia PA, USA. Catalog number: Plexiglas acrylic GS transparent UVD 2458). This creates an enclosed 301 * 31 * 32 cm volume (interior space, x * y * z; length * width * height) with a removable lid. To the backmost wall of the arena a 32mm hole is cut to allow the arena access tube to be connected to the arena. Paper wall panels with stripes of a 5cm periodicity are printed (full size pdfs for printing are provided in the repository associated with this publication [77], at: *protocol_downloads/custom_equipment_cad_files/Insect_flight_arena_with_frame/panel_5.03cmPwidth.pdf*) and cut to size to be attached to the arena’s long sides and bottom (3.02 by 0.32 meters) with masking tape. To the end walls a cut white paper sheet is stuck. Full details, itemized components and CAD files are available in [77], at: *protocol_downloads/custom_equipment_cad_files/Insect_flight_arena_with_frame/*within files *Arena_and_support_assembly.FCStd* and *Reference_for_Arena_and_support_assembly.docx*.
18. Support and camera mounting frame. Built to support the flight arena, cameras and AprilTag plates. Constructed from 40mm Aluminum profiles (Item Industrietechnik, Solingen, Germany, article number 0.0.026.33) and associated fittings for this profile system (Item Industrietechnik, Solingen, Germany). Notable design features are the presence of horizontal profiles above and below the arena support level for cameras, light arrays and AprilTag attachment. Full details, itemized components and CAD files are available in [77], at: *protocol_downloads/custom_equipment_cad_files/Insect_flight_arena_with_frame/*within files *Arena_and_support_assembly.FCStd* and *Reference_for_Arena_and_support_assembly.docx*.
19. Artificial flowers. Two types of artificial flower are used, at least one of each are required. Both utilize discs constructed from 3mm thick opaque yellow acrylic (Rohm and Haas, Philadelphia PA, USA. Catalog number: Plexiglas acrylic GS colored yellow opaque 1H01) cut to a 250mm diameter disc with a 32mm diameter hole in its center (using a Lazer cutter, see above for model). This hole allowed flowers to be attached to feeding reservoirs (see below) allowing bees to reach food within by going through the central hole in the artificial flower disk (artificial flower disc can be seen in figures S1-S3).

*Type a*. A falcon tube collar and wick design (similar to that used in [78]) This is constructed with the following parts as shown in figure S1: two 50mm sections of 32mm diameter acrylic tube (Grünke Acryl, Schwelm, Germany); a T-junction connector (Flowcolour, Zaozhuang Shandong, China, Item name: Acrylic PMMA fitting T-piece connector, diameter: 32mm 1”); a falcon tube (Corning Inc, Corning NY, USA, Item name: Falcon 50 mL High Clarity Conical Centrifuge Tubes) filled with sucrose solution, a cotton wool wick (made from a rolled cylinder, Paul Hartmann AG, Heidenheim Germany, item name: Hartmann Watte, article number: 4046871010515) and a 3d printed ‘collar’ (made using the same 3D printing materials listed above, and CAD files found in [77] at *protocol_downloads/custom_equipment_cad_files/3D_printed_parts/reinforced_collar.stl*). This collar screws onto falcon tubes, replacing the falcon tubes’ lid. In this ‘new lid’ is a 6.6 mm hole through which the wick is threaded. The top of this lid slots into the 32mm diameter acrylic tube. If the falcon tube were then filled with sucrose solution, the solution would wet the cotton wick. Bees accessing the tube would be able to feed from the wick. The wick then draws up more solution via capillary action as bees feed. This flower design sits level due to slotted stand cut to size from pieces of packing polystyrene. When empty of sucrose, the falcon tube and collar could be swapped out with fresh and full tubes or refilled.

*Type b*. a plastic jar design. The jar used was 6cm diameter, 3cm height with its lid removed and a sucrose-soaked wick (made as in type a) was placed inside. Artificial flower discs could be placed so the hole in the disk aligned with the open top of the jar (as shown in figure S2): allowing bees to access the wick by entering the jar through the hole in the disc.

1. Window-access. A 32mm diameter hole is cut into the lab window, to this an access tube for bees was constructed allowing bees to access the lab, and the feeders and arena within, by entering the hole in the flower disc. This window-access was constructed with the following parts as shown in figure S3: an artificial flower disc (as in item 19) sections of 32mm diameter clear acrylic pipe (Grünke Acryl, Schwelm, Germany), one 10cm, one 30cm and two 5cm lengths; an artificial flower disc (as in item 19) fixed to the end of the 10cm pipe with acrylic glue; two horizontal pipe connector fitting for 32mm tubing (Flowcolour, Zaozhuang Shandong, China, Item name: Acrylic PMMA fitting socket connector, diameter: 32mm 1”); a T-junction connector (Flowcolour, Zaozhuang Shandong, China, Item name: Acrylic PMMA fitting T-piece connector diameter: 32mm 1”); a falcon tube collar and wick (exactly as that used in *type a* artificial flowers). Access to the flight arena was initially blocked (as in figure S3), using Flowcolour, Zaozhuang Shandong, China, Item name: Acrylic PMMA fitting cap diameter: 32mm 1”). This allowed bees to learn to come to the lab window and expect rewards, but allowing the work within the lab discussed in steps A-I to be conducted without interference from bees. To this blocked arm, the arena access tunnel can be attached (see step J and K below).
2. Arena access tube. connecting the window-access to the flight arena. Made from the following constructed as in figure S4: 20mm (a length of 30cm and 50cm) and 32mm (one 10mm section) diameter clear acrylic pipes (Grünke Acryl, Schwelm, Germany), 2mm wide slits are cut along the width of the 50cm tube at 4cm intervals; two 45-degree pipe fittings for 20mm diameter tube (Flowcolour, Zaozhuang Shandong, China, Item name: Acrylic PMMA fitting 45-degree elbow connector, diameter: 32mm 1”); one horizontal pipe connector fitting (Flowcolour, Zaozhuang Shandong, China, Item name: Acrylic PMMA fitting socket connector, diameter: 32mm 1”) for 20mm tube and two horizontal pipe connector fitting for 32mm tube.
3. Arena access tube gates, made from garden plant labels (OBI Group Sourcing GmbH, Wermelskirchen, Germany, item name: Stecketiketten 12 cm, article number: 4007873253744). Used to control bee arena access.
4. Falcon tubes (Corning Inc, Corning NY, USA, Item name: Falcon 50 mL High Clarity Conical Centrifuge Tubes). For catching bees. If not able to release bees back outside shortly consider drilling air holes into the tubes.
5. Fly net (NHBS Ltd, Devon UK, item name: telescopic insect net, article number 245824). For catching bees.

### Software and datasets

The following software was used in the Braid tracking setup described here. The procedure for installation of software beyond the Linux Ubuntu operating system is described in the procedure section.

1. Linux Ubuntu 24.04 amd64 operating system for the PC running Strand-Braid. Installed exactly as described on the Ubuntu tutorial site: https://ubuntu.com/desktop/docs/en/latest/tutorial/install-ubuntu-desktop/. Some familiarity with using the Linux operating system and the Unix command line is useful.
2. ptpd (Linux Precision Time Protocol Daemon, Version: PTPD2 version 2.3.1), to implement Precision time protocol (PTP version 2 [79]) across the PC and cameras, for synchronization of frame timestamps between gig-E cameras.
3. pylon API and pylon Viewer (Basler, Ahrensburg, Germany. Version 7.3.0) to allow communication between Strand-Braid with Basler cameras and for setup configuration and troubleshooting with cameras (available here: https://www.baslerweb.com/en/downloads/software/?downloadCategory.values.label.data=pylon&softwareVersion.data=7.3.0).
4. Strand-Braid (Andrew Straw, Strawlab, version, 1.0.0-rc.6), available open source at https://github.com/strawlab/strand-braid/releases/tag/1.0.0-rc.6
5. BRAIDZ viewer (Andrew Straw, Strawlab) access online via a web browser here: https://braidz.strawlab.org/ (version used dated: Feb 5th 21:55:20 2024). Used for quick visualization of BRAIDZ files (with the ‘.braidz’ extension).
6. Python (The Python Software Foundation, version 3.12.9) and pip (version 25.0). Used for Rerun installation, analysis and plotting data. Use of Conda/Miniconda or similar for environments for Python is advised but is optional (miniconda version 25.1.1 was used).
7. Rerun-cli (version 0.22.1), hereafter ‘Rerun’. Used for detailed visualization of Braid live tracking, calibration and data.
8. Braid config file, available in [77] at */protocol_downloads/config_files_for_protocol/config.TOML*. This file is used to configure Braid to requirements, the available file tunes Braid for the setup used in this protocol.
9. Basler .pfs config file, used to set camera settings when Braid is launched to suit the setup used in the protocol, available in [77] at: */protocol_downloads/config_files_for_protocol/Basler_a2A1920_features.pfs*.
10. Retracking parameters config file, used to apply alternative tracking parameters to Braid’s offline tracking, available in [77] at */protocol_downloads/config_files_for_protocol/retrack_paras.toml*.

## Procedure

*Note: With regards to the timing of steps in the protocol. The nature of the Braid system and setup means pauses in work can be taken after (and indeed during) almost all steps with minimal issues*.

### **A.** Initial setup

1. Set up the dividers near the flight arena and support frame and place the worksurface behind it such that the worksurface is shielded from view by the dividers.
2. Place the PC and switch on the work surface.
3. Attach the lenses to the cameras.
4. Connect the Switch and PC.
a. Connect the switch to external internet via an ethernet cable.
b. Connect the switch to the PC using the 10G Gigabit ethernet cable.

*Warning: do not attach the PC to the external internet via another port on the PC. This will cause future behavior of the PC to differ from that we describe*.

5. Turn on the PC and log in.
6. Ensure the PC’s Gigabit Ethernet connection is enabled. Go to *PC Settings→Network* and activate the toggle corresponding to the Gigabit Ethernet to ‘on’. In the PC described above this is the ‘Aquantia Ethernet’ toggle.
7. Configure the switch to accept Jumbo Frames. This needs to be done on the PC network settings and on the switch itself.
a. Configure the PC network settings
i. Go to *PC Settings→Network* and then click on the wired network options cog button for the ‘*Aquantia Ethernet*’, this is to the right of the toggle clicked in step A6. This will open wired network options related to the switch in a new settings window.
ii. In this wired network options window navigate to the *Identity* tab and set the value of *MTU* to 9000 bytes.
b. Configure the switch itself via the switch’s own settings page, the *Instant On* portal.
i. In a browser enter the IP address of the switch. This will bring you to the switch settings portal.

*Note: the IP addresses of connected devices can be found using the command* arp -a *in the Linux terminal* [80].

ii. Enter the password for the switch.

*Note: If this is the first time you have logged into the switch’s settings portal you will need to use the default password (this will be provided with the switch, within the administrator guide or online:* https://arubanetworking.hpe.com/techdocs/AOS-CX/10.13/HTML/fundamentals_8400/Content/Chp_IniCfg/log-fir-tim-10.htm*). You will be required to create a new user and password for the switch when first logging in in this way*.

iii. Within the Instant On portal navigate on the left menu to the Switching tab and activate the Jumbo Frames toggle. Then click apply on the bottom right of the page. When this occurs you should receive a notification that this change needs to be saved and will only take effect after a device reboot.
iv. Save the changes to the switch settings by clicking the floppy disc save icon.

*Note: If you do not see this icon, click ‘Refresh’ or navigate back to the Dashboard tab (on the left menu). This may need to be performed a few times before the switch acknowledges settings have been altered and the save icon appears*.

v. Reboot the switch by turning it off and on. Once the switch has rebooted and web access restored, it is advised you return to the switch settings portal and the switching tab and verify these changes have been saved. If not repeat step A7b.

### **B.** Install and configure dependent software

1. Before beginning installation of software, in the Linux terminal run the following:

`sudo apt update`

and

`sudo apt upgrade`

*Note: all commands that require the super-user ‘sudo’ prefix will require inputting the user’s password in the terminal, assume this is needed each time ‘sudo’ appears in commands*.

2. Install the Basler pylon API and pylon Viewer.
a. Download the file ‘pylon_7.3.0.27189_linux-x86_64_debs.tar.gz’ from: https://www.baslerweb.com/en/downloads/software/?downloadCategory.values.label.data=pylon&softwareVersion.data*<u>=</u> <u>7.3.0</u>* at the linked site select the pylon 7.3.0 box in the downloads and then the “pylon 7.3.0 Camera Software Suite Linux x86 (64 Bit) - Debian Installer Package”. This will trigger the download.
b. Unzip the ‘tar.gz’ file downloaded (right click on the *tar.gz → Extract*)
c. Install pylon API and the associated Basler camera drivers using the following in a command line terminal launched in the extracted folder (either *right click on the open folder* → *Open in Terminal,* or open a terminal window directly and navigate to the folder with the `cd` command):

`sudo apt-get install ’./pylon_7.3.0.27189-deb0_amd64.deb’`
`’./codemeter_7.40.4997.501_amd64.deb’`

d. Once the commands from B2c have finished, restart your PC, and log back in.
e. Open pylon Viewer.

*Note: If pylon Viewer does not open, ensure steps B2a-d have been performed correctly, if so, consult troubleshooting problem 1*.

f. Connect all cameras to the Aruba switch with the ethernet cables. Within the *Devices* window of pylon Viewer each camera should appear listed under *GigE*.
g. Confirm the cameras are functional by opening each camera and selecting *Continuous Shot* (camera icon) from the top menu bar. You should see the cameras’ live feeds appear in a viewer tab as this is done.

*Note: If devices do not appear in the list of GigE cameras, or do not provide a live feed when Continuous Shot is selected consult troubleshooting and the Basler support:* https://www.baslerweb.com/en/learning-support/.

h. If all cameras function within pylon Viewer, you may now close pylon Viewer.

*Note: The Basler pylon API does not allow more than one piece of software to operate the cameras at once. Thus, fully close pylon Viewer after use, to allow Strand-Braid to operate the camera properly. Assume from now on that if pylon Viewer is used, it is closed after that step. See troubleshooting*.

3. Install ptpd using the following command:

`sudo apt install ptpd`

4. Configure the ptpd network. The ptpd’s configuration at download will not be configured to a specific PC and will not automatically activate. The ptpd configuration must be edited for Braid. Namely, with the PC running Braid as the ‘Master’ clock to which other ‘Servant’ devices on the network (the cameras) synchronize (see [79] for more details of PTP networks).
a. ptpd on installation saves a ‘minimal’ example config at the path ‘/usr/share/ptpd/ptpd2.conf.minimal’. Copy this config to ‘etc/ptpd2.conf’by running the following in the command line terminal:

`sudo cp /usr/share/ptpd/ptpd2.conf.minimal /etc/ptpd2.conf`

*Note: the file directory ‘etc’ can be accessed in the file navigator by going to ‘other locations → Ubuntu → etc’*

b. Identify the port interface ID for the Gigabit connection to the switch, hereafter the <PORT INTERFACE ID>, by either:
i. In the command terminal, enter:

`ip link | grep “state UP”`

This will display something like the following: “3: <PORT INTERFACE ID>: <BROADCAST,MULTICAST,UP,LOWER_UP> mtu 1500 qdisc mq state UP mode DEFAULT group default qlen 1000” Where <PORT INTERFACE ID> will be replaced by some sort of alpha-numerical code (e.g. “enp3s0”).

ii. Open the pylon IP Configurator (in *pylon Viewer →Tools → pylon IP Configurator,* or *Apps→ pylon IP Configurator*). This will display something like the window shown in figure 1 (be aware in figure 1 five cameras are connected to the switch. The *<PORT INTERFACE ID>* is the left aligned name entry after the green circuit board icon.
c. Edit the ptpd2.conf file (that was copied in step B4a) so the PTP network occurs over the Gigabit switch connection and uses the PC’s clock (as opposed to a camera) as the ‘Master’.
i. Enter the <PORT INTERFACE ID> for the Gigabit connection to the switch as the entry for ’ptpengine:interface’ (line: 10) like so:

ptpengine:interface=<PORT INTERFACE ID> that is, if the <PORT INTERFACE ID> were ‘enp3s0’ ptpengine:interface=enp3s0

ii. Edit ptpengine:preset (line 16) like so:

ptpengine:preset=masteronly

d. Edit the ptpd file at “/etc/default/ptpd” (note this is a different file from the one edited in step B4c) as follows:
i. on line 3: START_DAEMON=yes
ii. on line 6: PTPD_OPTS=”-c /etc/ptpd2.conf”

*Note: depending on the permissions of the user on your PC, either or both the above files may require the super user (sudo) to write and edit. To do this one can use the terminal editor nano, sudo nano <FILE path>, or edit a copy of a file and replacing the original using sudo cp (as we did in step B1)*.

e. Restart the PTP daemon with the following terminal command:

`systemctl restart ptpd`

f. Confirm the PTP network is running with the following terminal command:

`systemctl status ptpd`

This will produce an output similar to figure 2.

5. Install Rerun.

**Figure 1:**
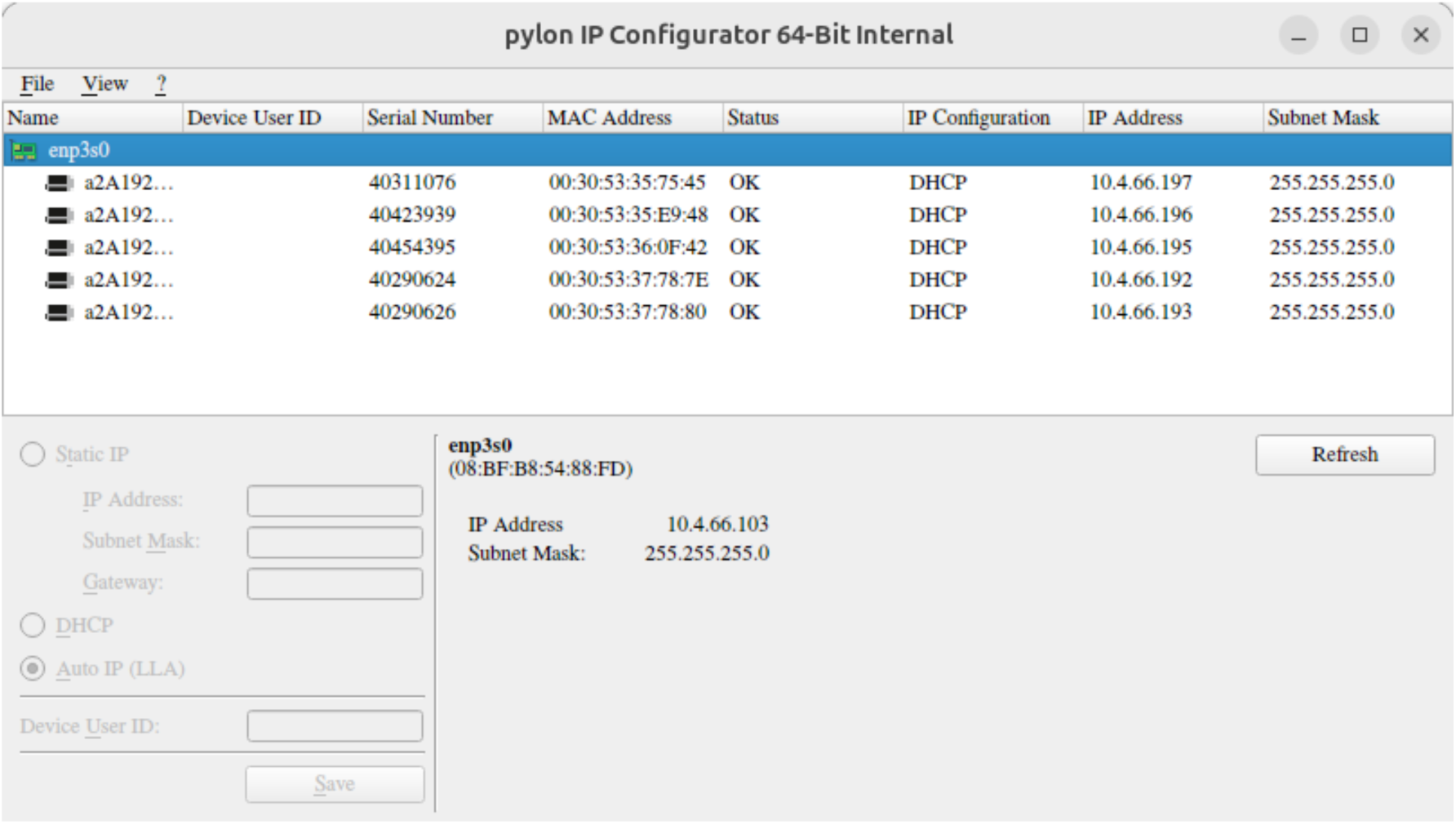
Basler’s pylon IP configurator, as it appears when 5 gigE cameras are connected to the switch. Note the green circuit board icon (on the selected line in the top left) and the <PORT INTERFACE ID> for the Gigabit connection to the switch here ‘enp3s0’.

**Figure 2:**
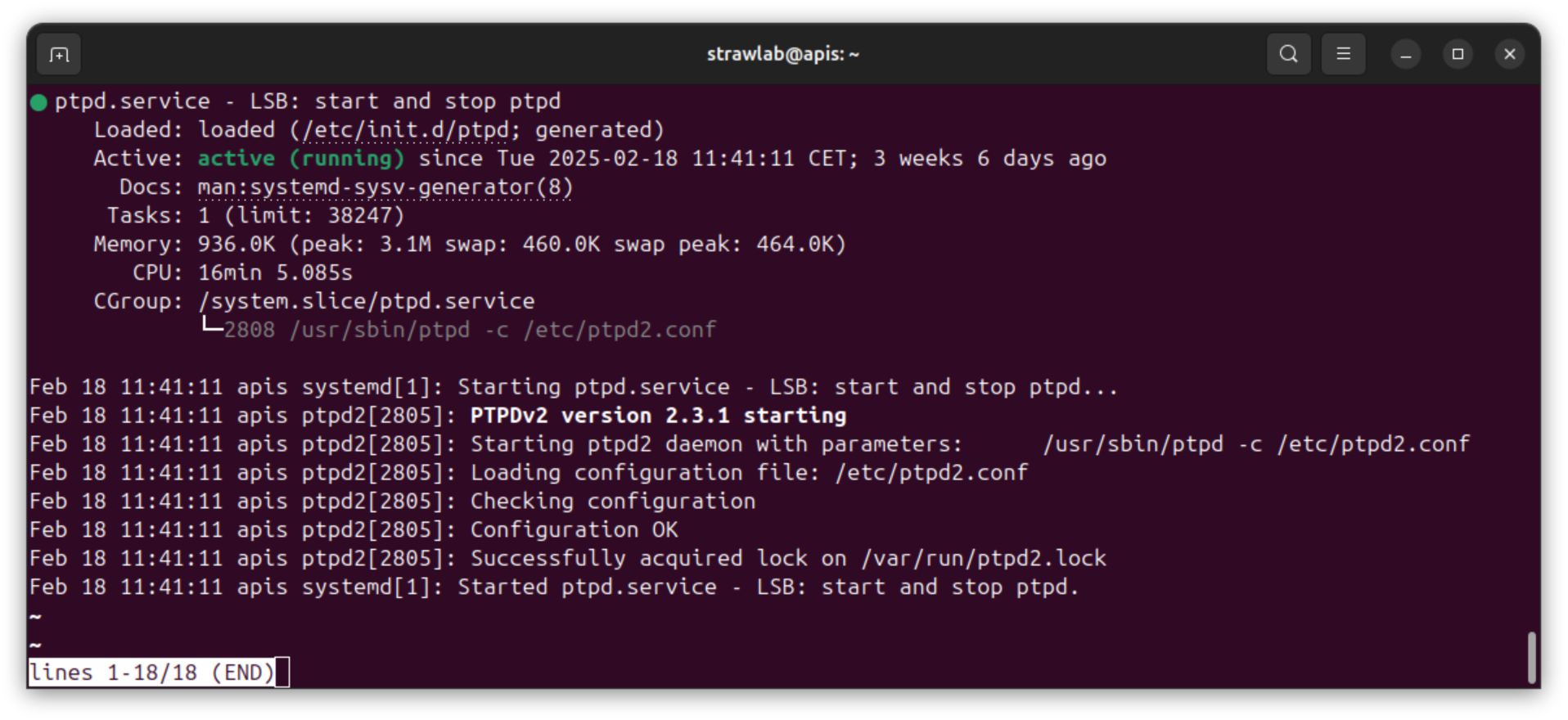
An example output of the terminal command *systemctl* status ptpd when ptpd2 is correctly configured. Appearance of a “Started ptpd.service” log entry indicates the ptpd network has correctly started. If this does not occur confirm step B4 has been correctly carried out.

*Note: Rerun can also be installed in different manners (see* https://rerun.io/*), we describe the simplest method using pip*.

a. Install pip. This can be done by either:
i. Using the following `sudo apt-get install python-pip`
ii. Using conda. Installation of miniconda as instructed in the following: https://www.anaconda.com/docs/getting-started/miniconda/main should create a base environment with pip installed.

*Note: If using conda all commands using the word ‘`rerun`’ must be run in the same conda environment. If your PC is being used solely for operating Braid it may be advisable to install Rerun in the base conda environment (see* https://www.anaconda.com/docs/main *for instruction on using conda environments)*.

b. Install Rerun using the following command

`pip install rerun-sdk`

c.Confirm the installation of Rerun by running the command

`rerun`

If installed correctly this will launch Rerun’s launch page. If this page opens, it can be closed for now.

### **C.** Install Strand-Braid

1. Download the github release of strand-braid. This protocol uses version 1.0.0.-rc.6 from https://github.com/strawlab/strand-braid/releases/tag/1.0.0-rc.6. Within the *Downloads* section of the release pages are ‘.zip’ files containing the Debian installer ‘.deb’ files for each supported Linux distribution (see figure 3). Click on *strand-braid-ubuntu-2404-1.0.0-rc.6.zip* to download the 2404 version.

*Warning: Strand Braid is supported on multiple distributions of Linux, for this protocol we use ubuntu 24.04 (although the protocol for Strand-Braid is similar across versions). Be careful that you select the correct version of strand-braid for the distribution (if using ubuntu 24.04 use 24.04 etc.). If the wrong distribution version of Braid is installed it may still function but may lead to performance issues or errors from certain functions*.

2. Unzip the Strand-Braid release ‘.zip’ file (*right click in file navigator → Extract*). Navigate to the unzipped folder. This will contain the ‘.deb’ installer file and a README.txt (which details installation of Strand-Braid).
3. Install Strand-Braid itself with the following in the command line in a terminal window opened in this location:

**Figure 3:**
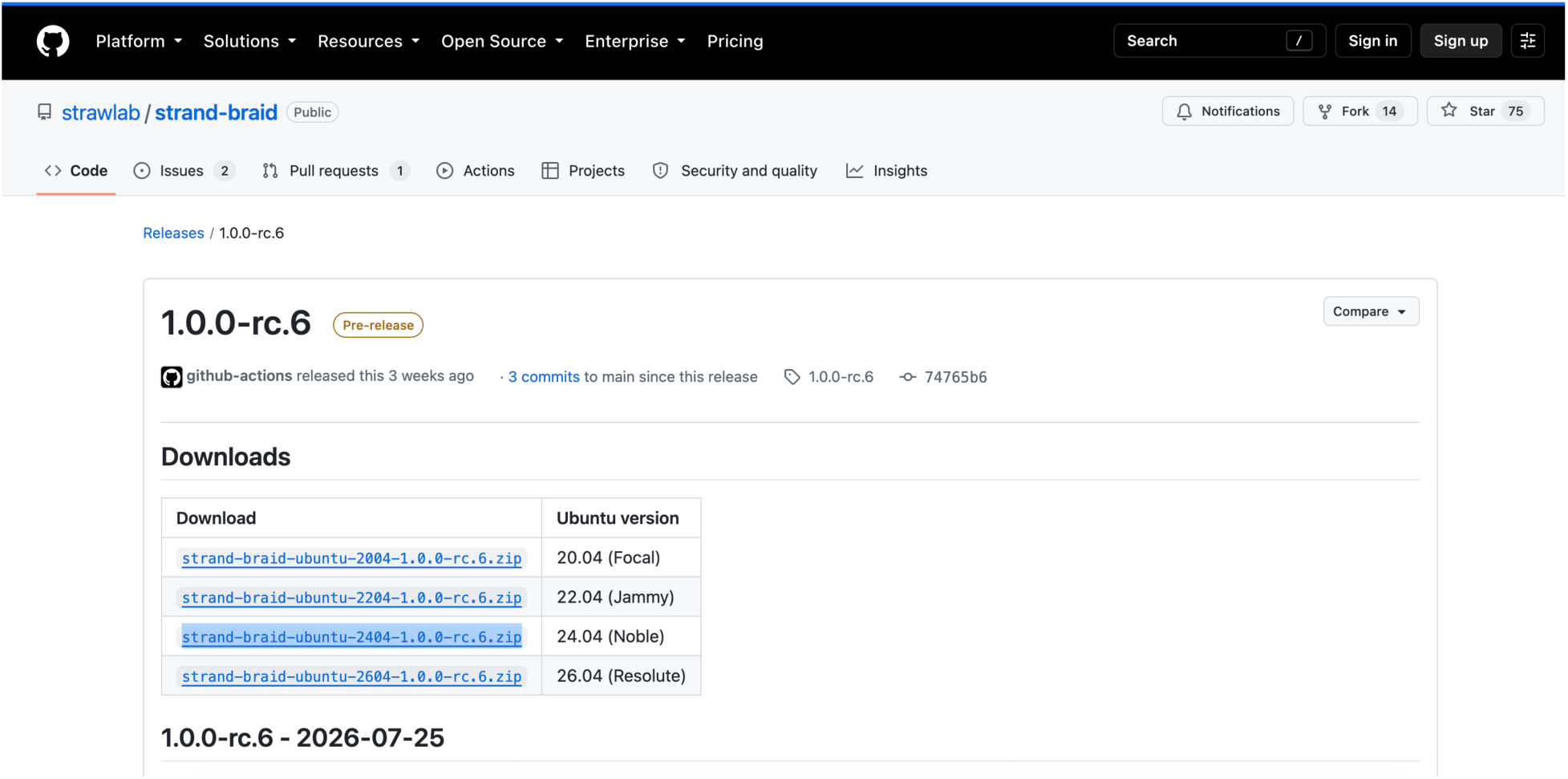
The Strand-Braid release on Github, note the *Downloads* tab and the release .zip files. Highlighted in blue is the 24.04 Linux version, clicking here will initiate download of the .zip file as indicated in step C1.

`sudo apt install ./strand-braid_1.0.0-rc.6-1_amd64.deb`

4. verify strand-braid’s installation by opening a new terminal window (anywhere) and running the following command:

`strand-cam --version`

if Strand-Braid installed correctly, this command should display the version information of Strand Camera.

*Note: Strand-Braid is supported on other distributions of Linux and will continue to receive support, and addition of features after this protocol. Other releases (including releases that predate and postdate this protocol) are available from* https://github.com/strawlab/strand-braid/releases*. Here releases are ordered newest to oldest. The procedure for installation of other releases of Strand-Braid will be like that described here. Specific instructions for each release are found in the README.txt file provided alongside ‘.deb’ files*.

### **D.** Launching Strand Camera on a single attached device

1. Attach a single camera to the switch using an ethernet cable.
2. run Strand Camera on this single camera with the following in a command terminal:

`strand-cam`

Strand Camera itself will launch with its default settings for the camera. Strand Camera’s graphical user interface (GUI) should launch in the PC’s default web-browser. Strand Camera itself will continue to run in the terminal it was launched in and provide an activity log (see Video 1).

*Note: If strand camera does not launch at this stage, verify the camera’s connection (with pylon Viewer, see step B2 and troubleshooting) and the strand-braid installation*.

3. Take note of the system camera name for the connected camera. This is needed to launch Strand Camera on specific cameras when multiple cameras are connected (step E) and to configure Braid (step F). Each system camera name is computed for each camera from its vendor and serial number. Thus, a Basler camera with the serial number ‘22441143’ would thus have the name Basler-22441143. The system camera name can be identified in 2 ways (both demonstrated in Video 1), either:
a. from the Strand Camera GUI. Where is given at the top of the page in the *Live view* tab, the title of which takes the form ‘Live view – <CAMERA_NAME>’.
b. from the activity log in the terminal Strand Camera was launched in. Where the following will appear in the log: ‘Camera “<CAMERA_NAME>” detected’.
4. Close Strand Camera by selecting the terminal the Strand Camera instance was launched in and either:
a. aborting the current task by pressing *Ctrl+c* while the terminal window running Strand Camera is selected.
b. closing the terminal window.
c. clicking on the ‘Quit Strand Camera’ button at the bottom of the GUI.

This will interrupt the GUI (as Strand-Cam is not running), the GUI browser window can also be closed (see video 1).

5. Disconnect the camera from the switch.

### **E.** Running specific instances of Strand Camera

1. Repeat step D for each camera. The command given in step D launches a single instance of Strand Camera on the first Basler camera detected by the system. Thus, it can only be used to launch Strand Camera on a specific camera if that is the only camera connected. To run Strand Camera on specific cameras when multiple cameras are connected, and later to configure Braid (see step F) one must obtain the system camera names for all cameras in the system.
2. Once all system camera names are obtained, connect all the Basler cameras to the switch.
3. Launch each camera on Strand Camera in turn using the following in the command terminal (demonstrated in Video 1):

`strand-cam --camera-name <CAMERA-name>`

where <CAMERA-name> is the system camera name of the camera. Thus, a camera with the name ‘Basler-22441143’ would be launched with:

`strand-cam --camera-name Basler-22441143`

*Note: to launch multiple instances of strand camera at once, you need to open multiple terminal windows (one for each camera)*.

4. Close all instances of Strand Camera (see step D4).

### **F.** Configuring and launching Braid

1. Create a Braid configuration toml for your system.
a. Download the config.toml from [77] at *protocol_downloads/config_files_for_protocol/config.TOML*. Each instance of Braid requires a configuration ‘.toml’ file [81]. Any options not specified within the configuration toml result in default values being used by Braid. The file config.toml has inputs required for running a 5 basler gigE cameras, synchronized at 40fps, with Strand Camera object detection and Braid mainbrain settings adjusted to suit tracking a bee within the arena volume.
b. Replace the system camera names in the config.toml file with those of your cameras obtained in steps D and E1. The entries for system camera names can be found on lines 26, 46, 65, 85, and 105 of the config.toml
c. Download the Basler ‘.pfs’ file from [77] at *protocol_downloads/config_files_for_protocol/Basler_a2A1920_features.pfs*, save this on your PC. We will refer to this file’s path as <CAMERA_SETTINGS_PATH> henceforth.
d. Edit the config.toml file so that the path indicated by each camera’s ‘camera_settings_filename’ is now <CAMERA_SETTINGS_PATH> the entries for each camera can be found on lines 27, 47, 66, 86, and 106 of the config.toml.

*Note: Different cameras can share a single camera settings file (as here) or can reference different camera settings files. New .pfs can be generated in pylon Viewer. Consult pylon Viewer’s documentation for more information:* https://docs.baslerweb.com/knowledge/saving-camera-features-or-user-sets-as-a-file-on-hard-disk.

e. Save the config.toml with your edits with the ‘.toml’ extension. It is advisable, but not necessary, to create a separate folder for Braid tomls in your ‘home’ directory. Henceforth the path to this toml file will be represented by <config.toml>.

*Note: In the config.toml provided, the mainbrain input ‘cal_fname’ (the calibration file path) is hashed out and therefore not read by Braid. We will restore this once the calibration is produced in step I. Braid will still launch without a calibration but will not be able to perform 3D tracking*.

*Note: The object detection parameters of Strand Camera and the braid tracking mainbrain parameters can be adjusted and there is a range of options. A full list of inputs and options for those inputs in Braid configuration files is available at* https://strawlab.org/strand-braid-api-docs/latest/braid_config_data/index.html [82]*. Additionally, the full list of settings given in the configuration toml will produce can be obtained by running:*

*`braid-show-config <config.toml>`*

2. Launch Braid using your configuration toml with the following command:

`braid run <config.toml>`

*Note: Braid will save certain files (AprilTag CSVs, MP4 video) in the current directory Braid is launched from*.

3. Allow Braid to launch. As with Strand Camera, Braid will continue to run in the terminal it was launched in and provide an activity log in that terminal window. If the system is installed and configured correctly Braid should launch similar to the instance shown in Video 2 .
a. Initially, Braid will provide confirmation in the activity log of the cameras it can detect and, if all cameras in the toml are present.
b. Braid will then attempt to synchronize these using PTP. While this is happening a series of activity log entries will discuss camera time offsets. Synchronizing the multiple GigE cameras may take a few seconds. Typically, synchronization takes longer the first time Braid is launched after cameras have been powered off. During this time, you may see the following warning in the terminal log: “launch time precedes device timestamp. Is time running backwards?” but this should stop appearing in the log after a few seconds (if it persists see troubleshooting problem 3).
c. When the Braid log reports ‘All expected cameras synchronized’ Braid is ready to be used.

*Note: If cameras do not synchronize, the Braid log will report ‘All expected cameras NOT synchronized’. Consult troubleshooting if this occurs*.

d. Access the Braid GUI by scrolling back through the Braid activity log until you find a link near a large QR code, click on that link or copy it to the browser (see Video 2).

*Note: the HTTP port for Braid’s control API (application programming interface) is set in the config toml using the input ‘*http_api_server_addr*’. In the example config.toml file provided this is set to the specific local port. This means each time Braid is launched it will be consistently accessible at the same web address in the browser. The QR code and link token provided can be used to access a running Braid instance by devices on the same network as the PC. This can be done by copying the link token into a browser or scanning the QR code with an appropriate device on the same network as the PC running Braid*.

4. Access individual Strand Camera instances that make up the Braid instance by clicking the camera names in the Braid GUI (Video 2) and navigating back to the Braid GUI using the *Back* buttons in the browser.
5. End the instance of Braid by:
a. selecting the terminal the Braid instance was launched in and aborting the current task with *Ctrl+c*.
b. selecting the terminal the Braid instance was launched in and closing the terminal window.
c. clicking the ‘Quit Braid’ button in the bottom of the GUI.

### **G.** Set up the tracking volume

*Note: In this protocol steps related to volume setup and calibration (steps G, H and I) are presented linearly. However, these steps represent parts of what is usually an iterative process. The accuracy of tracking achieved by latter steps is dependent on earlier ones and thus to achieve the desired level of accuracy going back to improve on earlier steps is often revealed to be necessary in latter steps. Furthermore, adjustments to improve accuracy of latter steps may also necessitate repetition of previous steps, see General notes 1 for more details*.

1. Attach a camera to a mounting plate then to a ball mount. It may be practical to temporarily disconnect the camera from the switch at this point.
2. Select a position for the camera and attach the mounted camera to the support frame at the selected position. When selecting position and viewing angle of a camera consider the following:
a. all points in the tracking volume should be viewed by as many cameras as possible.
b. collinearity and coplanarity of the camera’s positions (sharing a geometric line or plane respectively) should be minimized
c. camera fields of view should overlap as much as possible
d. cameras should be able to view common points about the volume to which AprilTag markers can be attached for extrinsic calibration (see figure 5 and step I).

The chosen positions for our tunnel volume are detailed in figure 4, and the resultant fields of view of each camera in the corresponding positions are detailed in figure 5.

3 .Attach the camera to an ethernet cable and launch strand-cam for the camera (as in step E3)
4. Observing the Live feed of the camera in the strand camera GUI. Adjust the viewing direction of the camera to view the arena.
5. When satisfied with the camera position, tighten the ball mount so that the cameras position is fixed.
6. prepare cameras lens settings for each camera.
a. Place the checkerboard, or another suitable target for focus, in the arena at a point in the camera’s field of view.
b. Viewing the live feed in Strand Camera, adjust focus, zoom and aperture of the lens so that the camera has the checkerboard in focus.
c. Fix the lens focus, zoom and aperture positions by tightening the screw heads on the lens.
7. Close the Strand Camera instance (as in D4).
8. Repeat steps G1-G7 for each camera, selecting a new camera position.
9. Attach AprilTags to points within your tracking volume. When placing tags consider the following:
a. Collinearity and coplanarity of tag plates should be minimized. Use the Sloped mounting blocks to support plates at non-flat angles.
b. Tags should be varied in xyz space (i.e. vertical and horizontal space).
c. Each tag should be detected by multiple cameras.

**Figure 4:**
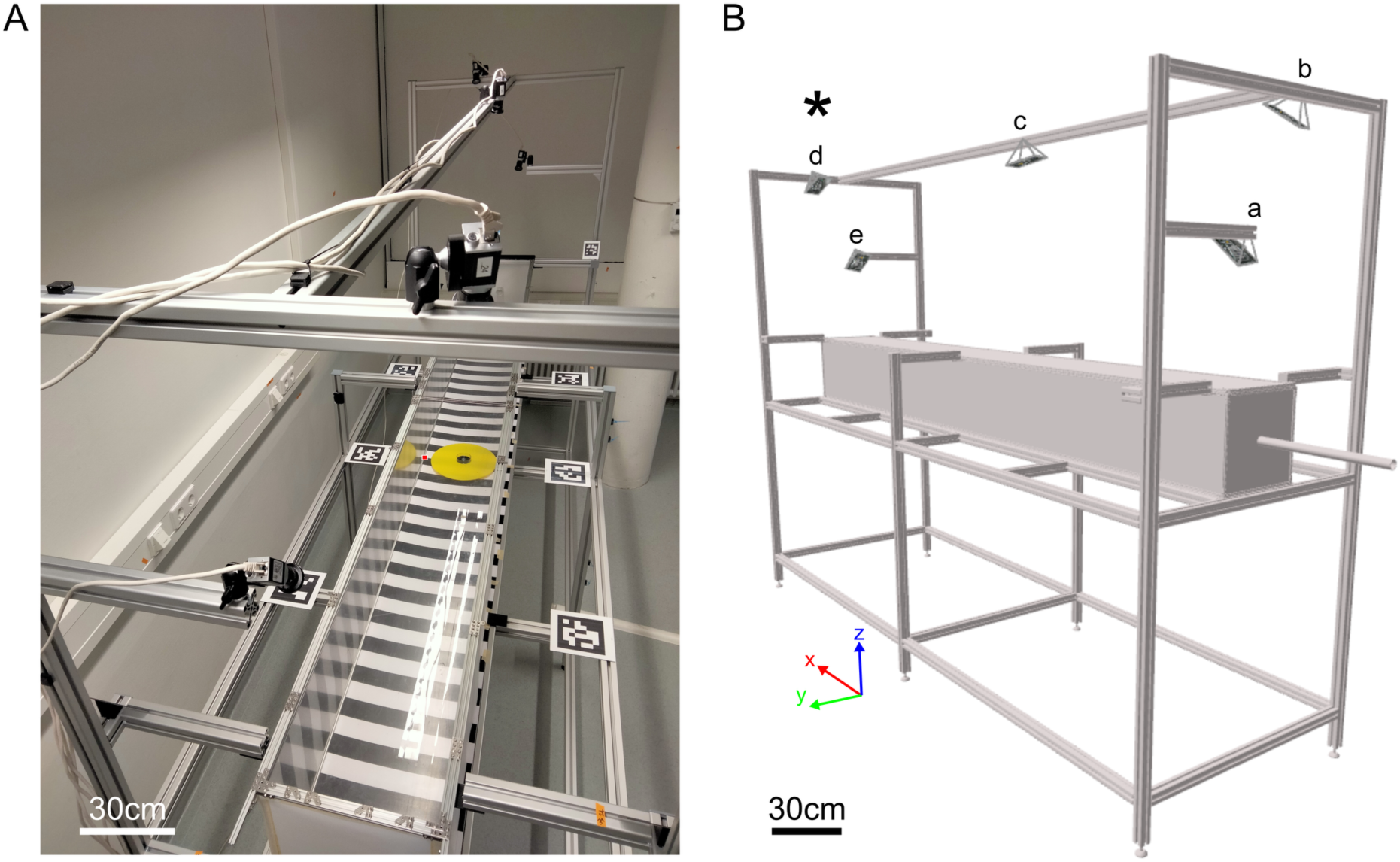
Example of suitable camera positions for 3D tracking of the flight arena tunnel with Braid. A) photograph of the positions of the cameras on the support frame. The red point symbol indicating the origin point of the coordinate system used to calibrate Braid (discussed in step G11). B) a 3D model of camera location and positions computed using the resultant extrinsic calibration (displayed using Rerun). Camera location (specifically the pinhole of each camera in the calibration model, which roughly corresponds with the camera sensor location) is indicated by pyramid points. Lowercase letter labels within panel B indicate camera identity with respect to the camera views in figure 5 (each letter corresponding with the panel label in figure 4; the view of the camera at position ‘a’ is seen in figure 5A, ‘b’ at figure 5B, etc.). The ‘✱’ symbol in Panel B indicates the position at which the photograph of panel A was taken. Axes indicated within panel B show the axes with respect to the coordinate system used in extrinsic calibrations (see step G11). Note how cameras vary such that they are non-coplanar and vary as much as possible in horizontal and vertical position without compromising view of the arena volume. Note that the 3D model does not show the camera mounts, hence cameras do not intersect with the frame. Note that the frame will subsequently have light arrays attached (step G12), these are not shown here. Note scale bars can only be an approximation due to image perspective view.

**Figure 5:**
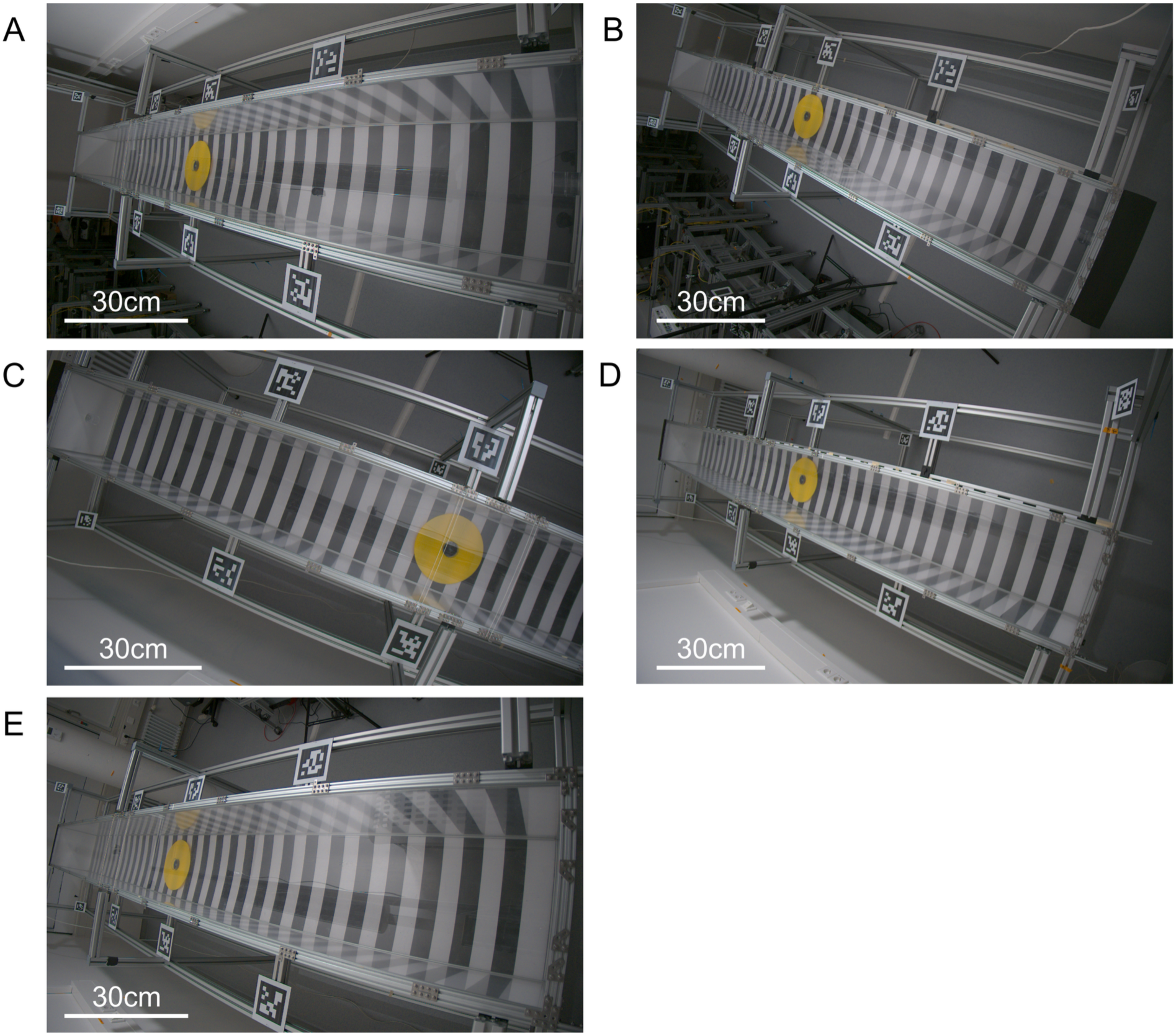
The fields of view of cameras in our setup, when the artificial flower is installed (described in step J). Each panel corresponds to a different camera’s view. Camera position is indicated in figure 4B (Panel A shows the view of the camera labeled ‘a’ in figure 4B, Panel B the camera labeled ‘b’ in figure 4B etc.). Note the cameras are positioned in a non-linear manner, some in line with the arena center, some aligned with the arena edges. Note that the cameras are also positioned in different viewing planes. The outermost cameras near the ends of the tunnel (A and E), view all the way down the length of the arena, as these are positioned lower than other cameras they view the arena at a nearer horizontal angle. The remaining cameras (B-D) each view the arena from above (nearer vertical). Note the highly overlapping fields of view and that most cameras (all except position C) can see the whole arena volume from a different angle. Additionally, note that AprilTags are visible around the arena by all cameras. Note scale bars can only be an approximation due to image perspective view.

The positions of our AprilTags (relative to the coordinate system described in figure 4) are given in [77] at *calibration_files/3d_coords_20250523.csv* and displayed within figure 5.

10. Verify AprilTag detections for each Camera.

*Note: AprilTag detection is computationally demanding. This may result in the Error described (with various solutions) in Troubleshooting Problem 2*.

a. Access an individual camera’s Strand Camera instance, either via Braid (step F4) or Strand Camera directly (step E).
b. In the Strand Camera GUI, navigate to and deselect *Object detection → enable object detection*, to turn off object detection.
c. Select the box at *AprilTag detection → enable detection*. If tags are detectable, the location of center of each tag will be highlighted with a green circle in the live view tab.
d. Looking at the *live feed* from the camera verify it can detect at least three AprilTags. If so, for each camera, continue to step G11. If not consider one of the following adjustments until this is the case for all cameras:

i. Adjust the position of the AprilTags.
ii. Adjusts the viewing angle of the camera.
iii. Adjust the position of the camera
iv. Adjust the focus of the camera.

*Note: while ‘at least’ 3 detections per camera are required, ideally more should be detected. In the system we demonstrate each camera detects at least 6 tags. It is advisable to maximize the number of tags each camera can see*.

*Note: AprilTags that are viewed at shallow angles, toward the more distorted periphery of the camera field of view, or appear small in the camera’s field of view may not be detected by the camera on every frame. This may mean the green detection point flickers in the ‘live view tab’. Such detections are still suitable for calibration*.

1. For each AprilTag measure its position in 3D space. Save these measured positions, into a CSV file with the columns: id,x,y,z. Where id is the AprilTag identity number. x,y,z. are the coordinates of the tag center in meters in the respective axes. An example of the CSV file documenting the locations of our tags is available in [77] at *calibration_files/3d_coords_20250523.csv*. This file and the path to it will be henceforth referred to as <3DCOORDS>.

*Note: The coordinate system used here will define values for data collected with the Braid system. Therefore, it is best for the x, y and z axes and their intercept position align with the volume. For our volume the origin point (x,y,z = 0,0,0) is the point on the inner right-hand side of wall at the base of the arena center (the position of the red marker on figure 4A). The x, y and z axes run parallel with the arena’s length, width and height respectively. These axes directions are marked on figure 4B*.

*Warning: Later processes of Braid assume a right-handed coordinate system* [as in 83,84]*, as is the case for the system described above. Use of a left-handed coordinate system will result in outputs being presented as mirrored*.

12. Attach lighting arrays to the support and camera mounting frame using the ball mounts. Angle the lights such that specular (mirror like) reflection from these lights in the arena is minimized. We find that arranging the lights so each faces along the arena’s length at a 45-degree angle from the top corners of the support frame works best for illuminating the arena. However, the ideal placement will depend on conditions of your setup (room wall color, extant light fittings etc.) and the exact placement of your cameras. The goal is to provide even illumination of the volume.
13. Connect lighting arrays to the mains. Circuit diagrams of lighting arrays and a rough sketch of lighting positions used are given in figures S5 and S6.

### **H.** Perform Intrinsic Calibrations (Checkerboard calibration)

*Note: checkerboard detection is computationally demanding. This may result in the Error described (with various solutions) in Troubleshooting Problem 2*.

1. Access an individual camera’s Strand Camera instance, either via Braid (step F) or Strand Camera directly (step E).
2. Activate checkerboard calibration for a camera.
a. In the Strand Camera GUI, turn off object detection as done in step G10b.
b. at *Checkerboard Calibration → Input: Checkerboard Size* enter the size of your checkerboard. For this purpose, the ‘size of a checkerboard’ is measured in the number of inner corners. That is, an 8 square by 8 square chessboard would have 7 internal corners in both height and width. If using the checkerboards we describe, enter 8 in the ‘width’ and 6 in the ‘height’ fields.
c. Select the box at *Checkerboard Calibration → Enable checkerboard calibration*.

*Note: If you wish to save the images collected during the checkerboard calibration and the checkerboard detection information, select the box at Checkerboard Calibration → Save debug information. The file path where debug information is saved will be indicated next to this box. By default a ‘.tmp’ (temporary) folder, thus it is advisable to save any debug information you wish to keep elsewhere once it is collected*.

3. Collect checkerboard detections.
a. The goal is to show the checkerboard to the camera so that Strand Camera can calculate the distortion to the image based on the apparent divergence of straight and flat lines of the checkerboard in the image. To get a good calibration (with mean reprojection error <1 pixel) we need to collect multiple images of the checkerboard that between them provide views of the checkerboard that:
i. Provide a good coverage of the camera’s field of view.
ii. View the checkerboard at flat and foreshortened angles.
iii. View the checkerboard at close and farther distances from the camera.

We list a series of target positions for the checkerboard that we aim to collect in table 1. This process is shown in Video 3.

**Table 1:**
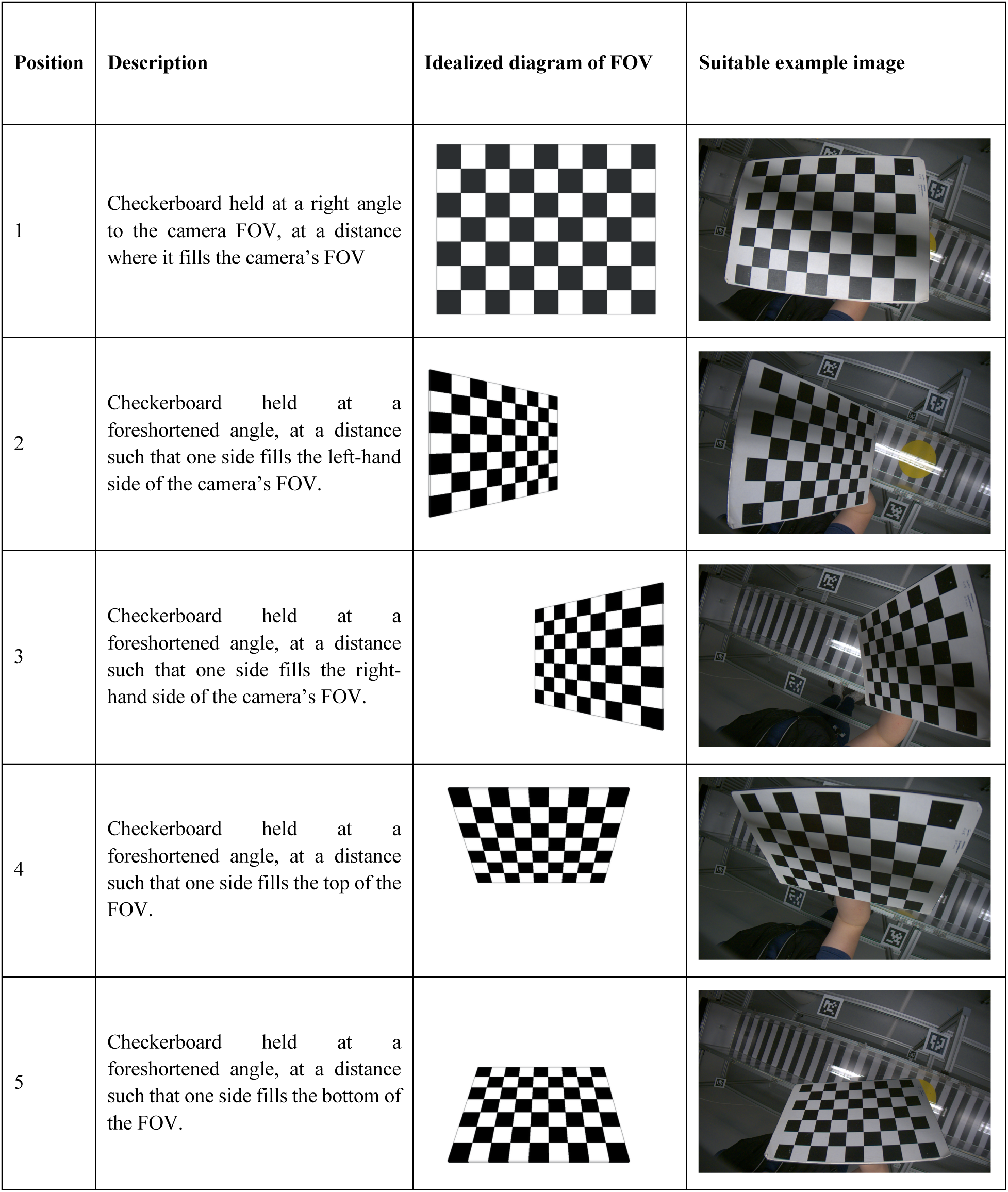
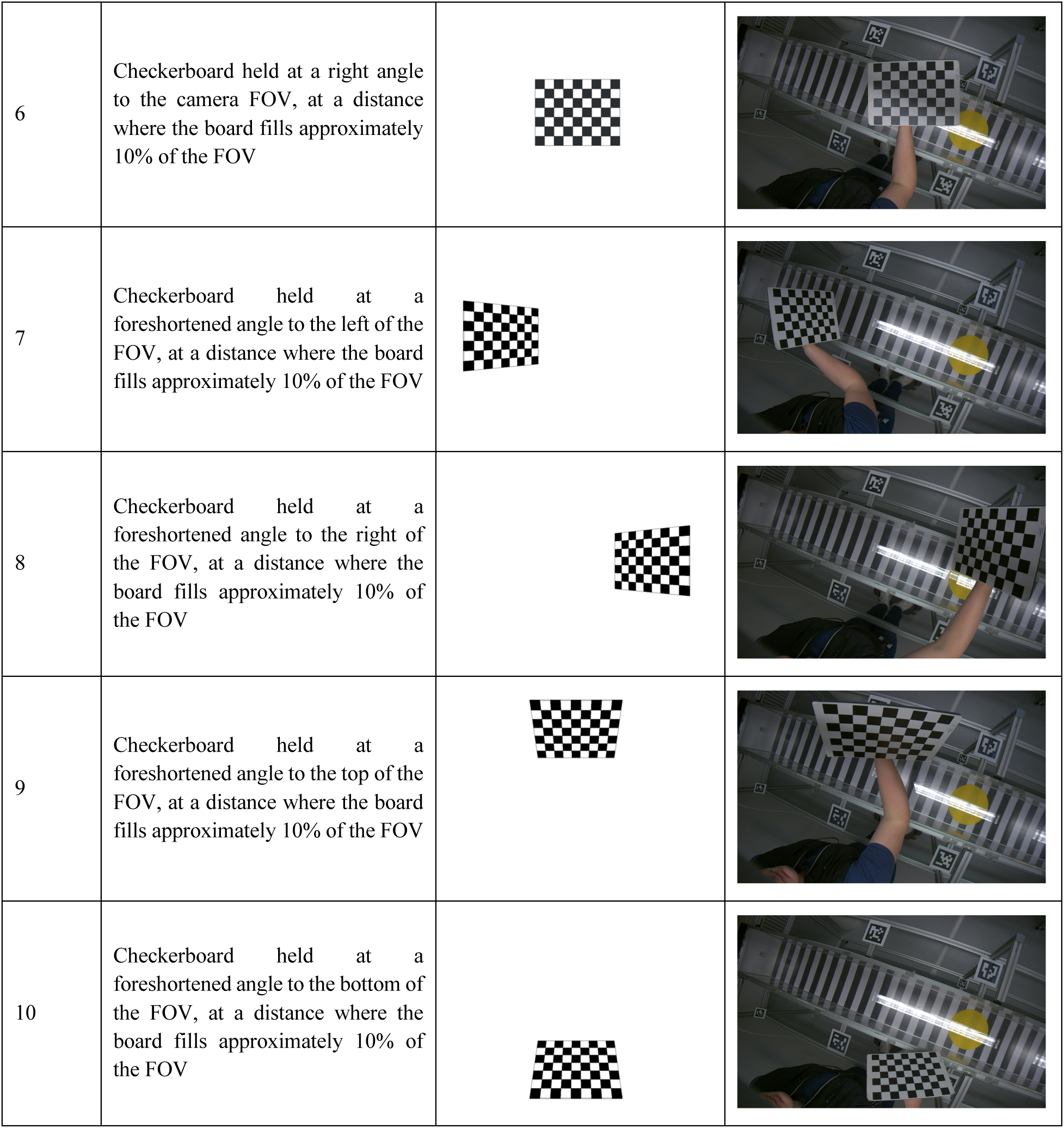
The target images we aim to collect during a checkerboard calibration. For each of our target views we provide: a description of view we wish to collect (see discussion in step H3); an idealized diagram of how the image we aim to obtain would look in the camera’s field of view (FOV, indicated by the blue box in idealized diagrams); and an example image from Video 3 which we consider a suitable replication of this view on a real camera. Note how distortion of the image is present in the example images but not idealized diagrams.

*Note: What is described here, is a process we have found consistently results in suitable checkerboard images for intrinsic calibrations in Braid. However different approaches can be successful. It is important to note that some approaches, particularly those that collect only images of checkerboard far away from the camera, can result in calibrations with unrealistic distortion estimates but low reprojection error. This may cause issues later in extrinsic calibration*.

b. Show the checkerboard to the camera so that you replicate a position listed in table 1.

*Note: Take care to not bend the checkerboards while performing intrinsic calibrations. Calculation of distortion assumes that the checkerboard’s lines are straight and flat and deviation in how these lines appear to the camera are the result of camera’s intrinsic features. Bending the board will thus introduce deviation in the appearance of the checkerboard that is not due to the camera. We advise holding the checkerboard by the handle attached to the back as shown in table 1 and video 3*.

c. Once the checkerboard is in the position desired wait and hold the checkerboard in that position until it is detected by Strand Camera. At *Checkerboard Calibration → Number of checkerboards collected* is a counter. Once a frame containing a checkerboard detection is made this number will count up. When this happens, the detected corners will be indicated in the live view with green rings. However, this can occur very quickly on the frame detected, so we advise looking at the counter.
d. Once the checkerboard is detected move the checkerboard smoothly to the next position listed in table 1, ideally keeping the checkerboard in the camera’s field of view.

*Note: Depending on exact camera positions and available space, you may not be able to assuming all positions in table 1 on each camera with the 30mm checked checkerboard. If so, use the smaller 10mm checkerboard for these positions*.

e. Repeat steps H3b-d. for each position listed in table 1.
f. Once checkerboards have been collected remove the checkerboard from the camera’s view.
g. Return to the PC and observe the number of checkerboards listed at *Checkerboard Calibration → Number of checkerboards collected.* This should read at least 10 checkerboards detected. However, it is likely that Strand Camera detected checkerboards collected while moving between the 10 positions. The goal is to have approximately 20-40 images, these should include those watched for in the 10 positions indicated in table 1. If there are many more, production of the calibration will take a long time; too few, and the calibration will likely be of low quality. If you have too many or too few checkerboards you may wish to perform a new checkerboard calibration.

*Note: You may find it difficult to quickly move between positions, this may have the result of many checkerboards being collected in each position. If so, consider reducing the frame rate of the camera (see General notes 2) until you can comfortably move between positions with a detection or two at each*.

h. If you wish to repeat the checkerboard calibration and collect checkerboards again. Press the button at *Checkerboard Calibration → Clear checkerboards*. You may then restart.
4. Produce the intrinsic calibration using the checkerboard images collected on step H3.
a. Press the button at *Checkerboard Calibration→Perform and save calibration* in the Strand camera GUI. This will produce an intrinsic calibration for the camera, [system camera name].yaml, it will also produce a copy of this calibration with a timestamped name [system camera name].[date]_[time].yaml. Both these files will be added to ‘home/.config/strand-cam/camera_info’.

*Note: Strand-braid uses the intrinsic calibration matching the camera name, the [system camera name].yaml file. Producing a new intrinsic calibration later overwrites this file. However, using the timestamped copy one can return to an old calibration by replacing the content of the file Braid uses with that of the timestamped copy*.

b. Deselect the box at *Checkerboard Calibration→ Enable checkerboard calibration* (and the box at *Checkerboard Calibration→ Save debug information* if selected previously).
5. Validate the intrinsic calibration.
a. Navigate to ‘home/.config/strand-cam/camera_info’ in the file explorer.

*Note: ‘.config’ is a ‘hidden file’. To view hidden files in the file navigator, click on the three bars symbol in a file navigator window ‘≡’ and ‘show hidden files’*.

b. Open the camera’s intrinsic calibration file and view the ‘Mean reprojection distance’ for the calibration (line 2 of the file). Confirm it is <1 pixel. If not, then repeat steps H1-5 for the camera.
6. repeat steps H1-5 for every camera.

*Warning: Camera intrinsic calibrations are specific to the physical lens adjustment of a specific lens and camera. If lens focus or zoom is changed after this point, the intrinsic calibration of the camera must be repeated. See General Notes 2*.

### I. Perform Extrinsic calibration (AprilTag position detection)

1. Collect AprilTag detection data for the extrinsic calibration.
a. Access an individual camera’s Strand Camera instance, either via Braid (step F4) or Strand Camera directly (step E), deactivate object detection (as in step G10b) and activate AprilTag detection (as in step G10c).
b. Activate AprilTag position recording by clicking on the circular record icon within the *AprilTag detection* tab. This will record a CSV in the directory from which you launched Braid.
c. Allow AprilTag recording to run for ten seconds. This will allow time for inconsistent detections (see notes following step G10) to be detected.
d. End AprilTag recording by clicking the square stop recording button (which replaced the record button clicked to start) in the AprilTag detection tab.
2. Repeat step I1 for each camera.
3. Move the unzipped CSV files to a single folder, the path to this folder will be henceforth referred to as <2D-dir>.
4. generate the extrinsic calibration by running the following in the Linux terminal:

`braid-april-cal-cli --apriltags-3d-fiducial-coords <3DCOORDS>`

`--apriltags-2d-detections-dir <2D-dir>`

`--intrinsics-yaml-dir <INTRINSICS>`

`--bundle-adjustment --bundle-adjustment-world-points-remain-fixed`

`--output-xml <output.xml>`

`--rerun-save <rerun_save.rrd>`

where <3DCOORDS>, <2D-dir> are the paths described as in steps G and I3.

<INTRINSICS> is the path to the folder containing the intrinsic calibration yaml files for each camera (collected in step H) normally this is ‘home/.config/strand-cam/camera_info’. <output.xml> is the path where the calibration will be saved, this path must end in .xml. <rerun_save.rrd> is a path to where a Rerun ‘.rrd’ file for visualizing the calibration will be saved, this path must end in ‘.rrd’. Braid can perform an optimization procedure, bundle adjustment, on all parameters within the calibration (see [85] for more information on bundle adjustment). By default, Braid does not alter camera intrinsic parameters, furthermore, inclusion of the additional argument ‘-- bundle-adjustment-world-points-remain-fixed’ orders braid to not perform adjustment on the 3D coordinates of the AprilTags. Thus, here bundle adjustment is restricted to extrinsic camera parameters.

*Note: The bundle adjustment procedure used by Braid can be altered or removed. Run* `braid-april-cal-cli --help` *for more details on available options. However, we advise (at least initially) using the restrictions detailed above*.

5. Validate the extrinsic calibration summary.
a. Observe the calibration summary provided in the output from the command run in step I4 or in the initial lines of the calibration ‘.xml’ file itself (saved at <output.xml>). This output has two main parts: the ‘Results from SQPnP algorithm using prior intrinsics’ and ‘Results after refinement with bundle adjustment model’.
b. Navigate to the section ‘Results after refinement with bundle adjustment model’ and inspect the following sections, verify if the following is the case:
i. In the section ‘3d distance between original and updated point locations’, verify for each tag ID the values read ‘0’.
ii. In the section ‘3d distance between original and updated camera center locations’, verify the adjustments are small, ideally below 0.05m.
iii. In the section “Camera parameters”, for each camera the ‘t_’, transverse, and ‘r_’, rotation, in the x, y, and z axes in the current calibration is given. Additionally, the intrinsic distortion parameters (fx, fy, cx, cy, k1, k2, k3, p1, p2) are also given for each camera, however as these have not been adjusted (due to restrictions applied to bundle adjustment, see above) these can be ignored. Verify that values of each camera’s transverse and rotation these appear to correspond with the cameras positions within the coordinate system you used.
iv. In the section ‘reprojection distance’ a table reports various reprojection distances (errors) in pixels, of each AprilTag visible to each camera. Additionally, the mean reprojection distance of each tag over all cameras that can detect it (rightmost ‘mean’ column) and of all tags each camera can detect (bottom ‘mean’ row). Lastly the mean reprojection error of all tags across all cameras is given, the mean reprojection error across the whole system, is provided (the intersection of the mean column and row). Low reprojection error is better (discussed in the section Validation of protocol). Verify that the mean reprojection error of all cameras is less than 10 pixels. Confirm that the calibration’s mean reprojection error (intersection of row mean and column mean) is small, ideally less than 5 pixels.

*Note: the above assumes that the calibration was generated with bundle adjustment settings as in step I4, if bundle adjustment settings are altered how outputs appear, and the acceptable values for a calibration, will differ*.

c. If at any part of I5b, you cannot verify values meet the criteria given, you must generate a new extrinsic calibration.
i. Confirm step I has been correctly performed up to this point. If not, repeat step I up to this point.
ii. If step I has been correctly performed up to this point, identify which tag(s) or camera(s) contribute to the high error. This will be indicated by higher reprojection distances corresponding to them in the ‘reprojection distance’ table.
iii. If a tag is associated with higher reprojection distance, consider remeasuring the position of that tag, or moving the tag so that it is detected by more cameras. Remember to update the <3DCOORDS> CSV file when remeasuring or moving tags before repeating step I.
iv. If a camera is associated with higher reprojection distance, consider adjusting the camera position, view or focus and/or performing a new intrinsic calibration (see sections G and H).
v. Once adjustments are made repeat step I and reevaluate the new calibration.
6. Visualize the calibration in Rerun.
a. run the following command:

rerun <rerun_save.rrd>

where <rerun_save.rrd> is as in step I4. This will launch the Rerun visualization of the calibration.

b. Within Rerun you will be presented with a 3D reconstruction of the calibration, using the coordinate system for your setup. Here numbered tags will be positioned at the center of the corresponding AprilTag and pyramid shapes indicate the location of the cameras. Additionally for each camera a reconstruction of its field of view, with AprilTags marked with corresponding numbered tags will also be presented. Using the time series scroller in the lower part of the image one can scroll from the beginning to end of bundle adjustment (if carried out). Looking at the Rerun reconstructions, particularly that at the end of bundle adjustment, confirm the arrangement of cameras and tags appears true to life. (For more details on using Rerun to visualize data see https://rerun.io/docs/getting-started/what-is-rerun).
7. Set your Braid config file to use the calibration.
a. Close Rerun and Braid, as detailed in step F .
b. Edit the Braid config file produced in step F to use the calibration.
i. Open <config.toml> in the Linux text editor.
ii. Delete the hashtag at the start of line 3, corresponding to the input ‘cal_fname’. Braid will now read this line when launched.
iii. Replace the path listed in the toml for ‘cal_fname’ with the calibration file produced in step I4. The line should look something like this:

cal_fname=”<output.xml>”

where <output.xml> is the path to the braid calibration file.

c. Save and close <config.toml>.
8. Restart Braid with the edited config.toml (as in step F). Now the Braid GUI should report the presence of a calibration file (see 00:37 on video 2). At this point Braid is ready for 3D tracking.

*Warning: if a camera position (angle, translation, rotation) or the intrinsic state of a camera is altered necessitating a new intrinsic calibration, extrinsic calibration must be repeated. See General Notes 2*.

### J [optional]. Provide bees with experience of the flight arena

*Note: Bees lacking experience of the flight arena often do not fly straight to the artificial flower when they first enter the arena, even if they have experience visiting artificial flowers or the window-access likely because they lack experience of the arena (similar observations are described in [76]). Allowing bees to experience the arena before tracking lead bees to have directed searches of the arena [73–76]. It is, however, possible to track a completely naïve bee within the arena and not conduct step J*.

*Warning: opening the arena and replacing and refilling the artificial flower feeding reservoirs while bees have access to the arena risks bees escaping into the lab and should be conducted by someone experienced in handling bees in such a setting. If bees escape, catch them in falcon tubes and fly net and release them outdoors*.

1. Place an artificial flower attached to a feeding reservoir (type a) in the center of the flight arena.
2. Connect the arena access tube, without gates inserted, to one of the T-junction arms of the window-access to allow bees visiting the window-access free access to the flight arena (see figures S3 an S4).
3. Maintain the rewards in the arena daily as follows:
a. Remove the artificial flower and feeding reservoir from the flight arena.
b. Replace the reservoirs in the artificial flower and window-access with new tubes filled with sucrose and replace the wick if there is visible mold or a change in color. Dispose of the used wicks.
c. Return the artificial flower and feeding reservoir to its place in the arena.

*Note: Due to bee foraging habits it is safest to conduct the feeding reservoir refills at night, when bees are less active and likely to be attempting to access used feeder reservoirs while they are being replaced with fresh ones*.

4. Continue step J3 until bees have learnt the location. This will be evident by bees’ flight to the artificial flower and the presence of bees within the flower feeding reservoir tubes.

*Note: the exact time step J3 takes will depend on many factors including the weather. We allowed bees free access to the arena for ten days*.

5. Once bees have learnt, clear the arena of bees.
a. Place gates within the arena access tube and allow bees leaving the arena to pass by lifting the gate as the bees approaches. Prevent bees trying to enter the arena by leaving the gates down in front of them.
b. Once bees are no longer leaving the arena. Remove the lid of the arena.
c. Using the fly net and falcon tubes capture any remaining bees and release them outdoors.
d. Remove the artificial flower from the flight arena.

*Hint: Unplugging the lights prior to removing the lid of the flight arena can be helpful as the dark conditions will discourage bees from flying making their capture easier*.

e. Using a paper towel damp with ethanol cleaning solution wipe down the inside of the arena.
f. Return the arena lid.

### **K.** Collecting 3d tracking data

1. Prepare the arena for tracking.
a. Place an artificial flower with a jar feeding reservoir (type b) in the center of the flight arena. Ours was placed such that the center of the flower hole was at -0.043, 0.17, 0.03 x, y, z coordinates, within the system described in step G11.
b. Place the gates into the arena access tube (if not already there).
c. Attach the arena access tube to the window-access (if not already attached).
2. Launch Braid and access the GUI (as in step F).

3 [optional]. If saving video MP4 recordings alongside 3d tracking data, set the MP4 video recording frame rates. By default the MP4 recordings by Braid are ‘unlimited’ and records video at the maximum frame rate (40 FPS in our config.toml). However, recording high frame rate video alongside tracking data can overtax the PC-switch connection and lead to frames being lost and tracking interruptions. It is advised to set the video recording frame rates.

a. Access a camera’s strand camera instance as in step F.
b. Select the frame rate for the camera using the buttons at *MP4 Recording options→ MP4 Max Framerate*.
c. Repeat steps a and b for each camera. We have found the system described here can tolerate 4 cameras set at 40FPS and one set at 1FPS. However, to minimize the chance of tracking being interrupted, we typically set 4 cameras at 30FPS and one at 1FPS. When finished return to the Braid GUI.

*Note: adjusting MP4 recording frame rate has no effect on the recording frame rate of 3D tracking data. To adjust 3D tracking data recording frame rate, either lower the rate of the Braid system’s trigger pulse (see general notes) or select a frame rate for each camera in the camera’s strand camera GUI→ Object Detection →CSV Max Rate*.

4. Lifting the gates as a bee within the access tube approaches, allow a bee (the ‘focal bee’ for this recording) through the gates in the access tube closing these gates behind the focal bee. Isolate the focal bee in a tube between two gates in the tube. Ensure there are at least 2 closed gates between the focal bee and the window-access. This will prevent other bees in the window-access reaching the arena while tracking in the arena.
5. Allow the focal bee the ability to access the arena by removing the gates in the access tube between it and the arena.
6. **Immediately** return to the PC behind the divider and click the circle record icon at *Record .braidz file* in the Braid GUI to begin recording positional data. Ideally this should be done before the focal bee enters the arena while the bee is walking through the arena access tube.
7. [optional] If MP4 video is required press the circle record icon at *Record .MP4 files* in the Braid GUI <u>after</u> BRAIDZ file recording has been started.

*Hint: to save the hard drive space taken up by video you may wait until the bee has entered the arena. We can assess when this is via the braid GUI by viewing a strand camera instance or the model live feed (see step L below)*.

8. Record the focal bee in the arena.
9. [optional] To end MP4 recording, click the square icon at *Record .MP4 files* in the Braid GUI <u>before</u> ending BRAIDZ recording (step K10). Saved MP4 files will have names corresponding to the time and date of the start of recording and the system camera name. These videos will be saved in the directory the Braid instance was launched from.
10. To end the recording of the BRAIDZ file, click the square stop icon at *Record .braidz file* in the Braid GUI. Ending a BRAIDZ recording will result in a BRAIDZ file (more details in Data Analysis section) saved in a new directory *Home/BRAID-DATA*. This BRAIDZ file will have a name corresponding to the date and time of the start of recording and the file extension ‘.braidz’.

*Important: To ensure error free processing, particularly when exporting tracking to Rerun, recording of BRAIDZ files should begin before recording of MP4 files and end after the end of MP4 files, as described above*.

11. Once finished with a recording either allow the focal bee to return to the arena access tube opening gates for it, as in steps K4, allowing it to leave, or remove the lid to the flight arena, capture it and release it outdoors.

### **L.** Visualizing Braid tracking in real time

There are 3 ways to visualize the tracking state of the bee while collecting data:

1. You can access live feeds of the individual Strand Camera instances (as in step F) during recording and view the *Live feed* window. Here 2D detections will be indicated by a green ring about the object detected.
2. On the Braid GUI clicking *Model server*. This will lead you to the live feed of the detections providing you with the current object ID and a current tracking estimate.

*Note: Braid will assign objects it tracks separate object ID numbers. This object ID number will remain with an object while 3D tracking is maintained. Braid’s utilization of a Kalman filter to make 3D tracking estimates means brief loss of 2D detections can be accounted for. However, if the object remains completely still for a long period of time, the tracking of that object will be lost. This is because 2D detection make use of a model of the background image of each camera. By default, this is continually updating, thus a stationary object will begin to be incorporated into this background model. Similarly, if the target is obscured for a prolonged period, the tracking will be lost. Within the arena bees may be obscured for long periods when a bee goes under the artificial flower or the edges of the tunnel. Upon redetection, tracking will resume with a new object ID. Braid tracking settings controlling background model updating and how readily the object is discarded can be altered within the config.toml (see* https://strawlab.org/strand-braid-api-docs/latest/braid_config_data/struct.BraidConfig.html *for more details)*.

3. Using a live feed sent to Rerun.
a. before tracking a focal bee (before step K5) set the global parameter

export RERUN_VIEWER_URL=” rerun+http://127.0.0.1:9876/proxy”

b. start Rerun by running the command

rerun

and leave the Rerun window open.

c. Track a bee as in step K.
d. While tracking the bee (during K8) return to the Rerun window, within the 3D visualization you should see the current tracking state within the 3D reconstruction.

### **M.** Visualize the BRAIDZ file

There are 2 ways to quickly visualize BRAIDZ files collected by Braid:

1. View a BRAIDZ in the BRAIDZ viewer.
a. Access the BRAIDZ viewer via an internet browser at https://braidz.strawlab.org/ or app if installed locally.
b. Drag and drop the BRAIDZ file to view into the viewer. This will present x and y detection values for each camera, then a pair of 3D reconstruction plots (x against y, and x against z).

*Note: if the BRAIDZ viewer is installed locally it can be set as the default application for BRAIDZ files. However, this is discouraged as it will replace Rerun viewer as the default*.

2. View a BRAIDZ 3d reconstruction in Rerun.
a. Double click on a BRAIDZ file. This will launch Rerun viewer. Rerun will present a 3D reconstruction of the 3D tracking data from the BRAIDZ file alongside 2D detections in frames for each camera. Rerun’s data visualizations can then be navigated by frame using the timeframe scroller in the lower half of the Rerun interface. Labels indicating the object ID corresponding to 2D detections and 3D tracking will also be attached to points within these views. When tracking an object, camera views can show up to two dots while tracking a single bee. One is the detection made by the strand camera instance for that camera, if such a detection occurs. The other is the reprojection of the object’s position in the 2D view based on Braid’s 3D estimate. This reprojection point is displayed by Rerun in the camera views even when the estimate is outside the camera’s field of view.
3. Visualize BRAIDZ data with video using Rerun. To do this a Rerun rrd file needs to be produced using the BRAIDZ and video files.
a. run the following command to produce a Rerun rrd file:

`braidz-export-rrd <PATH_TO_BRAIDZ> <PATHS_TO_VIDEO>`

where <PATH_TO_BRAIDZ> is the path to the BRAIDZ file, and <PATHS_TO_VIDEO> are the paths to each video collected simultaneously with that BRAIDZ file. Additional <PATHS_TO_VIDEO> arguments can be added for each camera, however the more <PATHS_TO_VIDEO> added the longer production of the rrd will take.

b. View the resultant rrd file by running the following:

`rerun <PATH to rrd>`

Video 4 shows a demonstration of Rerun’s 2D and 3D reconstructions. For more details on using Rerun see https://rerun.io/docs/getting-started/what-is-rerun.)

### **N.** Tracking multiple targets with Braid

1. Temporarily change the number of objects Braid tracks.
a. Launch Braid.
b. In Braid navigate to the Strand Camera instance of a camera (as in step F4) go to *Object Detection→ Edit configuration* input box and change the ‘max_num_points’ to the desired number of points (this number must be an integer, such as 1, 2, 3 etc.).
c. Press the ‘->’ arrow between the *Edit configuration* and *Current configuration* input boxes to send this change in object detection configuration to the camera
d. Confirm that the change is now present in *Object Detection→ Current configuration.* If not, repeat steps O1a-c or observe any error warnings below the ‘*Edit configuration*’ box. Here errors such as incompatible or incorrect inputs to Current configuration will be explained and advice is given.
e. Within the *Live view* tab of the Strand Camera instance you should observe more 2D detection indicators (green rings) when objects move within the camera field of view.
f. Repeat steps N1a to c on each camera. Once complete the Braid system is now ready to track multiple objects.
g. Repeat step K, but this time allow more than one focal bee (up to the number you have set `max_num_points` to) into the arena to be tracked.

*Note: adjusting the number of tracked objects in this way, while useful for quick checks, does not alter the configuration Braid launches into when restarted. If Braid is closed you must repeat this step. If you wish to keep changes made to Braid’s configuration you must edit the config.toml file as in step N2*.

2. Edit the config file from which Braid is launched. Adjusting the config file in this way allows Braid to launch with settings, such as ‘max_num_points’, set at the desired value.
a. open the <config.toml>
b. Edit the value under ‘max_num_points’ within the sections ‘cameras.point_detection_config’ for each camera to the desired number of points (again, this number must be an integer). This is found on lines 38, 58, 77, 97 and 117 for each camera.
c. save and close the config file.
d. once the above is complete Braid can be launched as in step F. This will result in Braid being launched with the edited configuration, being able to track the desired number of points. Within the *Live view* tab of the Strand Camera instance you should observe more 2D detection indicators (green rings) when objects move within the camera field of view.
f. Repeat step K, but this time allow more than one focal bee, (up to the number you have set ‘max_num_points’ to) into the arena to be tracked.
3. Visualize the data collected in steps N1 and N2 as done in step M.

*Note: other configuration parameters, can also be changed using the same process as that described in step N. For further details on camera object detection parameters:* https://strawlab.org/strand-braid-api-docs/latest/braid_types/struct.BraidCameraConfig.html *and Braid mainbrain parameters:* https://strawlab.org/strand-braid-api-docs/latest/braid_types/struct.TrackingParams.html.

### **O.** Retrack a BRAIDZ file

1. When visualizing 3D tracking data collected live, as in step M, you may identify areas where 3D tracking estimates are absent but 2D detections are made. Braid prioritizes producing 3D estimates at low latency. This means that if 2D detection information from cameras is delayed (due to being held in the switch memory, for example) and received out of order, Braid will not ‘wait’ for past frames, or detections from cameras that have not arrived. Braid will prioritize data for the current frames in calculation of 3D positions (the ‘Kalman estimates’). However, if tracking and production of BRAIDZ files continues without errors, the corresponding detection data is logged in the BRAIDZ file. Retracking allows Braid to recalculate the 3D positions using all the detection data collected for each frame.
2. retrack a BRAIDZ file using the following command:

`braid-offline-retrack --data-src <BRAIDZ> -o <OUTPUT>`

Were is the

<BRAIDZ> ‘old’ BRAIDZ file to be retracked and the

<OUTPUT> file path of the resultant retracked BRAIDZ , this must end in ‘.braidz’. This command will produce an output BRAIDZ with retracked data using the same calibration and tracking parameters as used as when the original live data was collected.

*Note: braid will not allow you to overwrite a BRAIDZ file using the above command so*

<BRAIDZ> *cannot be the same as*

<OUTPUT> *and you must change the output filename if performing multiple retracks*.

3. Visualize the new output BRAIDZ file as in step M. Depending on the BRAIDZ file, retracking may slightly improve quality of 3D tracking. Most often these improvements will include:
a. 3D position estimates may be adjusted or added for frames where they were absent in files collected live.
b. Trajectories of the same target that were previously assigned different BRAIDZ object IDs (as discussed in step L2) may be unified as a single object.
c. Low confidence projections when objects are lost may be reduced.
d. Braid object IDs within the BRAIDZ file will be altered to start from 0.
4. It can be beneficial to retrack data with altered Braid tracking parameters or a different calibration (such as one of a higher quality). Retracking with alternative configurations can be performed using:

`braid-offline-retrack --data-src`

`<BRAIDZ> --tracking-params <TRACKING-params_toml>`

`--new-calibration <NEW_CALIBRATION>`

`-o <OUTPUT>`

where

<BRAIDZ> and

<OUTPUT> are as in the command given in step O2 and <TRACKING-params_toml> is a ‘.toml’ file containing the new tracking parameters (an example of such a toml file is provided in [77] at *protocol_downloads/config_files_for_protocol/retrack_paras.toml*) and <NEW_CALIBRATION> a new calibration file produced as described in step I. --tracking-params and --new-calibration are optional inputs and either or both can be excluded (note that excluding both inputs is the command in step N2).

## Data Analysis

When you collect tracking data with Braid, it is saved in the BRAIDZ file format (with the extension ‘.braidz’, usually in a folder called ‘BRAID-DATA’ in the home directory). This file format is a zip file with specific contents pertaining to Braid tracking. The key component of a BRAIDZ file is the ‘Kalman_estimates’ datafile. This provides Kalman estimates of each tracked objects position in xyz space (in meters) and directional velocity (m/s) in x, y and z directions, both within the coordinate system used for calibration. It also provides uncertainty values of these estimates [9,86]. For more details of specific other BRAIDZ components see https://strawlab.github.io/strand-braid/braidz-files.html.

Within the strand-braid GitHub repository generic Python scripts and Jupyter notebooks for accessing data in .braidz files are provided, see https://github.com/strawlab/strand-braid/tree/main/docs/user-docs/analysis/. Additionally, alongside datasets provided with this publication, we provide the specific Jupyter notebooks for plotting data collected by Braid using this protocol and performing the further validation step conducted in the next section [77]. These provide examples of how to access braid data from BRAIDZ files in Python and how to generate 2D and 3D graphs (as in figure 6-8) of data collected with Braid. BRAIDZ files can also be unpacked using the `unzip` command [87] on the BRAIDZ , and then the `gunzip` command [88] on the content to view the raw CSV files. Using the raw ‘Kalman_estimates.csv’ datafile data collected with Braid can be analyzed in the platform of your choice.

**Figure 6:**
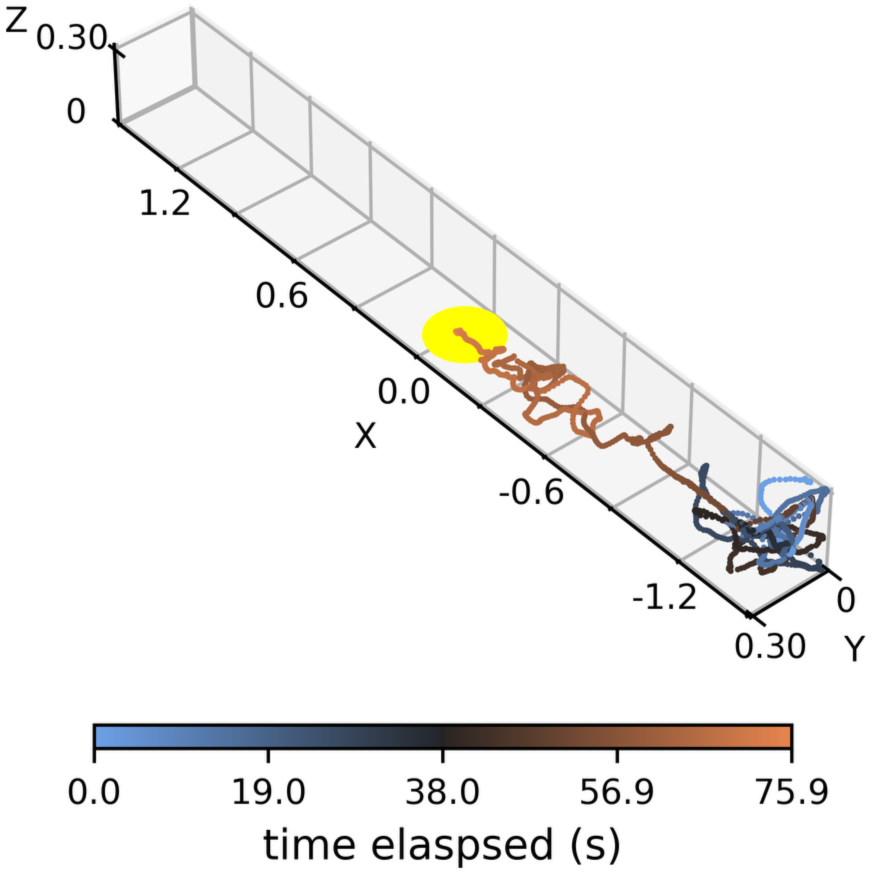
An example of a 3D plot of a single bee (the bee with the time identifier ‘1427’, see [77] for detail on bee ID numbers). Axes are represented in meters and describe the flight arena’s 3D axes. Points plotted are the bees’ position during tracking according to Braid. The yellow disk indicates the artificial flower position, note tracking terminates as the bee enters. Color of plotted points indicates time elapsed from when the bee enters the arena and is detected as an object by Braid. Code for generation of such graphs is available alongside datasets included with this publication.

## Validation of protocol

Braid’s 3D tracking is based on the same tracking algorithm as its predecessor software Flydra [9], thus the principles behind Braid’s 3D tracking algorithm are well validated. Flydra’s 3D tracking estimates have been validated against real life objects previously [9] and Flydra has been successfully used to assess and measure 3D positions of insects and other animals (e.g. [9–19]). Furthermore, versions of Braid that predate the release described here have also been used previously [1–8]. Additionally, within a meta-analysis of 3D tracking tools, Braid was appraised highly compared to alternative software [89] based on performance tracking insects, particularly mosquitoes.

Using the above protocol up to step M (including step J, where bees were allowed to access the arena for 10 days before tracking) we tracked flights of 28 individual bees alone within the arena (data available in datasets included with this publication). We tracked each bee until they located the flower and entered it, stopped searching, attempted to leave the arena. Additionally, tracking was ended after 5 minutes had elapsed if none of the previous criteria were met. Of the bees tracked, 12 of these were successful flights where the bee was able to locate the flower within 5 minutes. Tracking data for 3 example flights are provided in figure 7 (tracking results for all successful flights are provided in figure S7, and figure S8). Braid can effectively track insects live against complex backgrounds, here contrasting stripes. As discussed in step O, retracking data can further increase quality of tracking. Braid is also highly suitable for live 3D tracking applications that require low latency such as integration with closed loop virtual reality or position triggered applications [17,11,16,4]. Braid is also capable of tracking multiple objects at a time. Figure 8 demonstrates the results of tracking multiple bees (in this case 3) simultaneously after following the protocol up to step N (including step J), further examples of tracking multiple bees simultaneously are provided (as BRAIDZ files) in the datasets associated with this publication [77].

**Figure 7:**
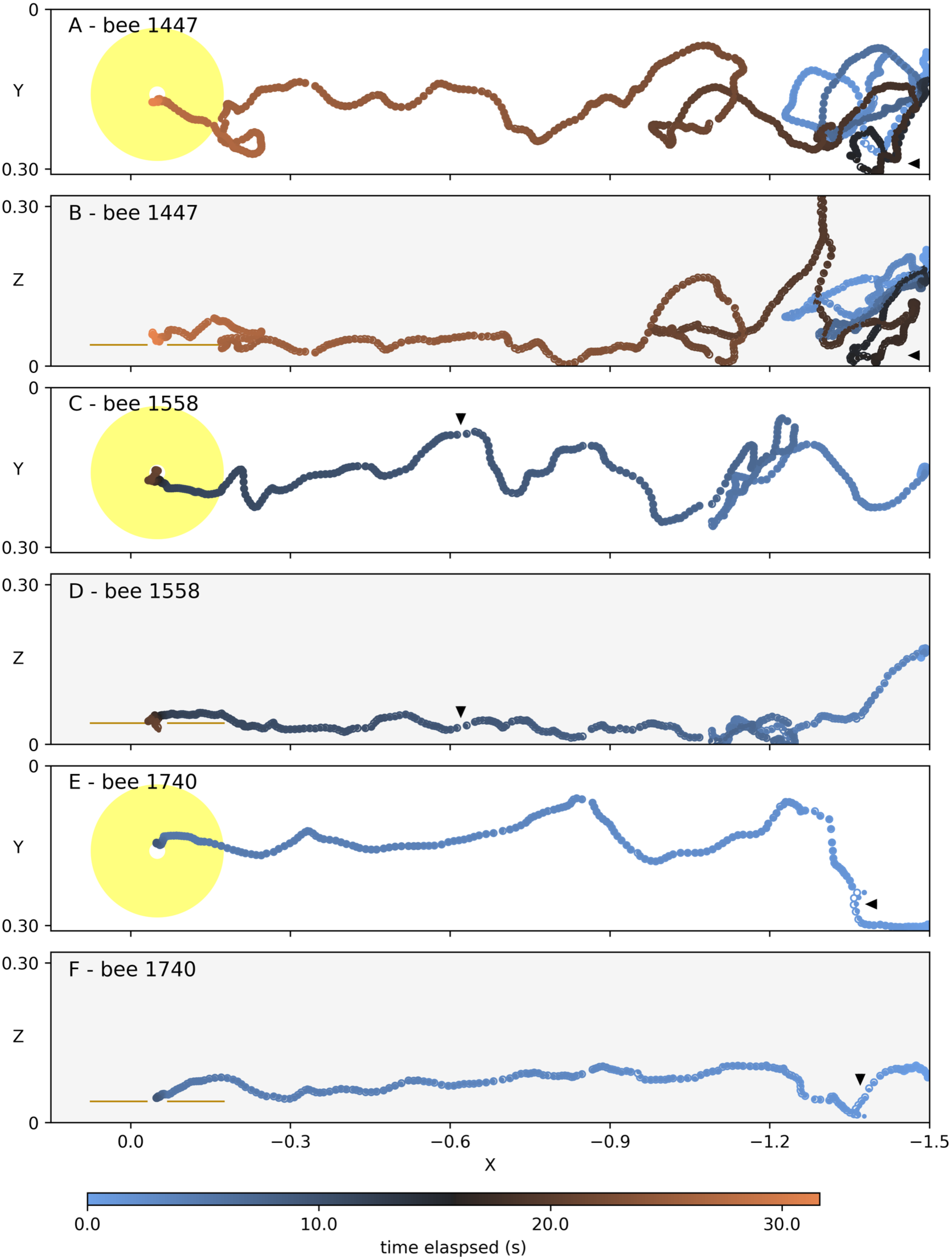
3D tracking results of 3 bees that successfully located the flower within the arena. Data is collected as described in the protocol up to step O (including optional step J). A pair of graphs are presented for each bee (individual bees are named by the time at which tracking began): A and B) bee 1447, C and D) bee 1558, E and F) bee 1740. Each bee’s trajectory plotted in x-y (A, C, E, G) and x-z (B, D, F, H) axes. Axes are in meters and are as described for the tracking arena. The artificial flower position during tracking is indicated by the yellow disc (xy graphs) or lines (xz graphs). Note that tracking ends when the bee enters the inside of the artificial flower. Smaller filled points indicate the tracking estimates obtained after data is retracked with the original tracking parameters from the config.toml provided (as in step O2). Larger white filled points indicate tracking estimates data retracked using the alternative tracking parameters provided in retrack_paras.toml but not changing calibration (as in step O4). Consequentially, a larger filled point indicates original tracking estimates correspond well with those with more constrained tracking estimates. Black triangles indicate a section in each trajectory where original and retracked Kalman estimates differ more noticeably. Color of point fill (original estimates) and edges (retracked estimates) indicates time elapsed (in seconds) since the bee entered the arena.

**Figure 8:**
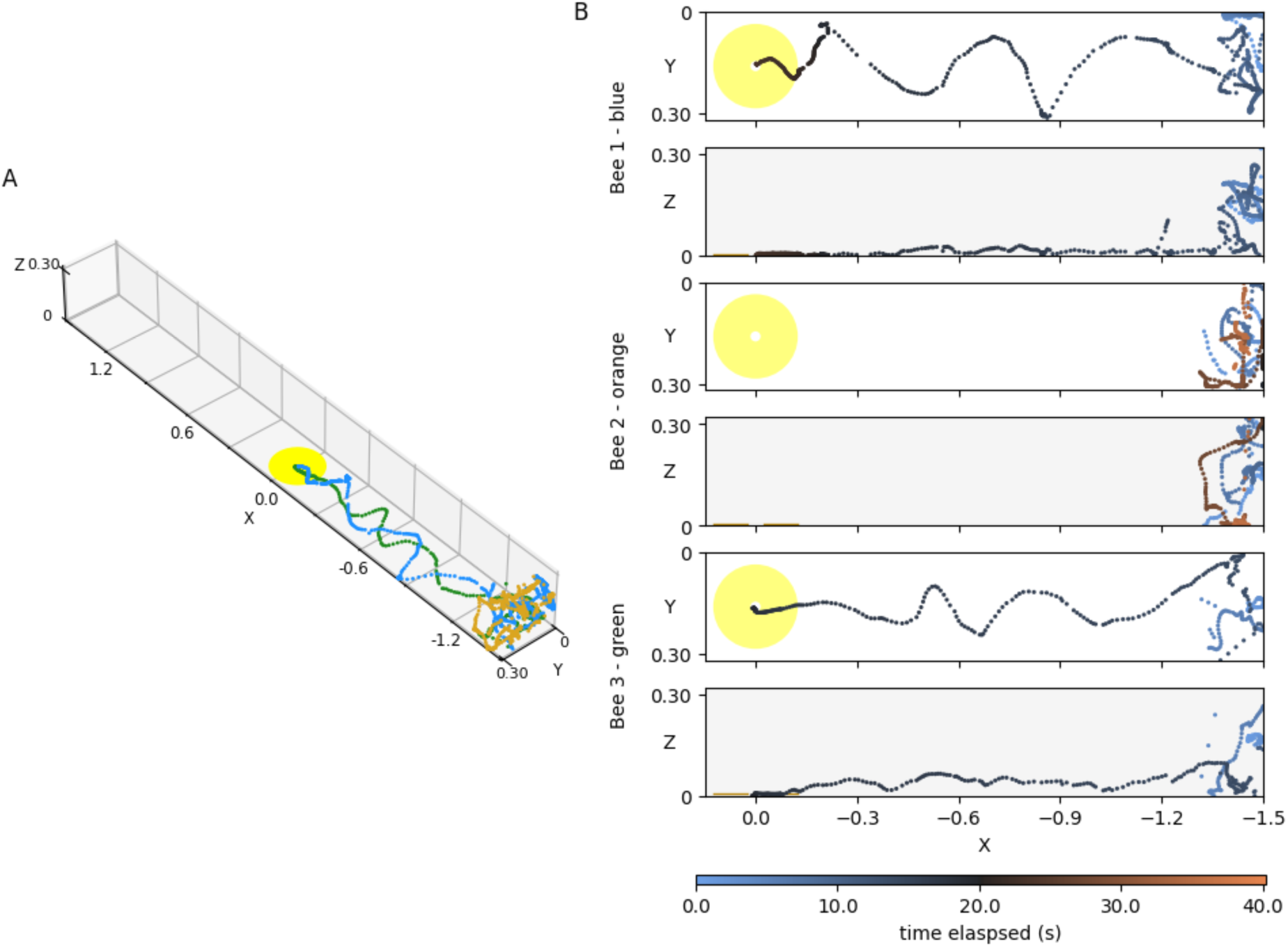
3D tracking of three bees simultaneously (specifically from the file *20251015_172023_retracked*.*braidz*, see [77]) within the flight arena. Data used is retracked data but maintaining the tracking parameters used in config.toml (as discussed in step O2). Tracking of the bees is plotted A) together rendered 3D and B) separately in 2D plots. This data was collected by following the protocol described above including the optional step J. Axes are represented in meters and describe the flight arena’s 3D axes. In A) color of points indicates individual bees, verified in Rerun. In B) for each bee a pair of plots are presented for each bee in the x-y (white background) x-z (grey background) axes. Each bee’s 2D plots can be identified in the 3D plot based on the label to the left side of x-y and x-z plots presented as the bee’s ID number and the corresponding color of that bee in panel A. In B) color of points indicates time elapsed (in seconds) since the first bee entered the arena. The artificial flower position during tracking is indicated by the yellow disc (A, and xy graphs of B) or lines (xz graphs of B). Other examples of simultaneous tracking of multiple bees are provided in the datasets associated with this publication [77].

During the procedure described above Braid conducts several validation steps on the calibrations it produces allowing evaluation of the system as it is prepared. These take the form of calculation of ‘reprojection distance’ also known as ‘reprojection error’. In general terms, reprojection distance is a measure of the quality of calibration models used by Braid. Reprojection is the act of re-plotting the location of a detected point using the estimate of that point’s predicted location derived from another coordinate system; this replotted point is the ‘reprojection’. Thus, reprojection error or reprojection distance, is the difference between the detected points location in the original coordinate system and that of the corresponding reprojection (see [85] for more details). Within Braid the ‘detected’ points are the 2D detections made by Strand Camera within each camera’s field of view, the reprojections are the corresponding points derived from calibration models (intrinsic and extrinsic). That is, the reprojections are estimates of where a location would appear on a 2D field of view given the calibration model. Within the Braid setup procedure this method is employed twice and reprojection distance always represented as pixels. We assess the quality of intrinsic calibrations in step H5 comparing the detected internal corners of the checkerboard against the camera’s intrinsic calibration (distortion) model estimate where checkerboard corners should appear. Second, we assess the quality of the 3D extrinsic calibration model in step I5 where, we compare the 2D detections of 3D points (the AprilTag positions) with the 2D reprojections of those positions derived using the 3D extrinsic calibration model. Using the protocol above one should be able to produce intrinsic calibrations with mean reprojection distances of <1 pixel. Those used to collect the data shown here <0.55 pixels (see camera calibrations provided in [77]). Using the protocol as described we were able to produce extrinsic calibrations with mean reprojection distance as low as 1.95 pixels (see [77] for examples).

To further validate the 3D tracking of the Braid setup described by this protocol we conducted a further validation testing the accuracy of Braid’s measurements compared to real life values. An LED wand with two small 2mm wide LED lights (Broadcom, Palo Alto CA, USA, article number HSMW-C170-U0000) was constructed using Makerbeam aluminum profiles (figure 9a). The distance between the centers of these LED lights was measured, using calipers, to be 0.2924m apart from each other. The Braid system was then set to track two points at once. To facilitate detection of LED lights the room lights were turned off and the arena walls and floor were covered by a matt black fabric sheet. The first of these measures ensured the background model for each camera was an even black image, so ensured the LEDs were the most different pixels viewed by the cameras and were tracked. The sheet, reduced reflections from the Perspex arena interfering with LED point estimates. The LED wand’s lights were turned on while in the arena and moved about the arena volume, while doing so the wand was raised and lowered and twisted back and forth within the arena. The combined speed of this LED motion (based on Braid’s Kalman estimates of velocity) was 0.25 ±0.11 m/s (mean ±standard deviation). Once the wand had been moved throughout the arena and back, the LEDs were turned off. Meanwhile 3D point estimates were collected by Braid. This BRAIDZ file was collected at 20 fps (see general notes 1) to reduce the processing demands and reduce the quantity of data collected while the wand was stationary as it was turned on and off. Video images were also collected. Mean reprojection error of the Braid calibration used was 1.95 pixels. Using Rerun (as in step M2) it was confirmed that the LEDs, were tracked throughout the sequence of motion. At time points where Braid detected two points within the arena the 3D Kalman estimates of both points were extracted from the BRAIDZ file and the distance between them calculated. Using this approach, we collected 696 estimates of a known distance (0.2924m) throughout the tracking volume. The distance estimates between the LEDs by Braid were accurate to the sub mm level and varied little, with the mean estimate of the distance being 0.2927m ±0.001 (mean ±standard deviation) and repeatability coefficient [90] of 0.00280m (figure 9). We thus find Braid to be highly accurate in position estimation, accurately and consistently estimating the LEDs to be within millimeters of their true position, allowing accurate estimates of distance between them. This is especially the case as much of this variation could be explained by how LEDs are 1.5mm wide and where the brightest spot on the LED appears to a given camera may be influenced by viewing angle. Thus, we might expect estimates of each LED position to vary by approximately 0.75mm even when Braid is accurately detecting the position of the LEDs.

**Figure 9:**
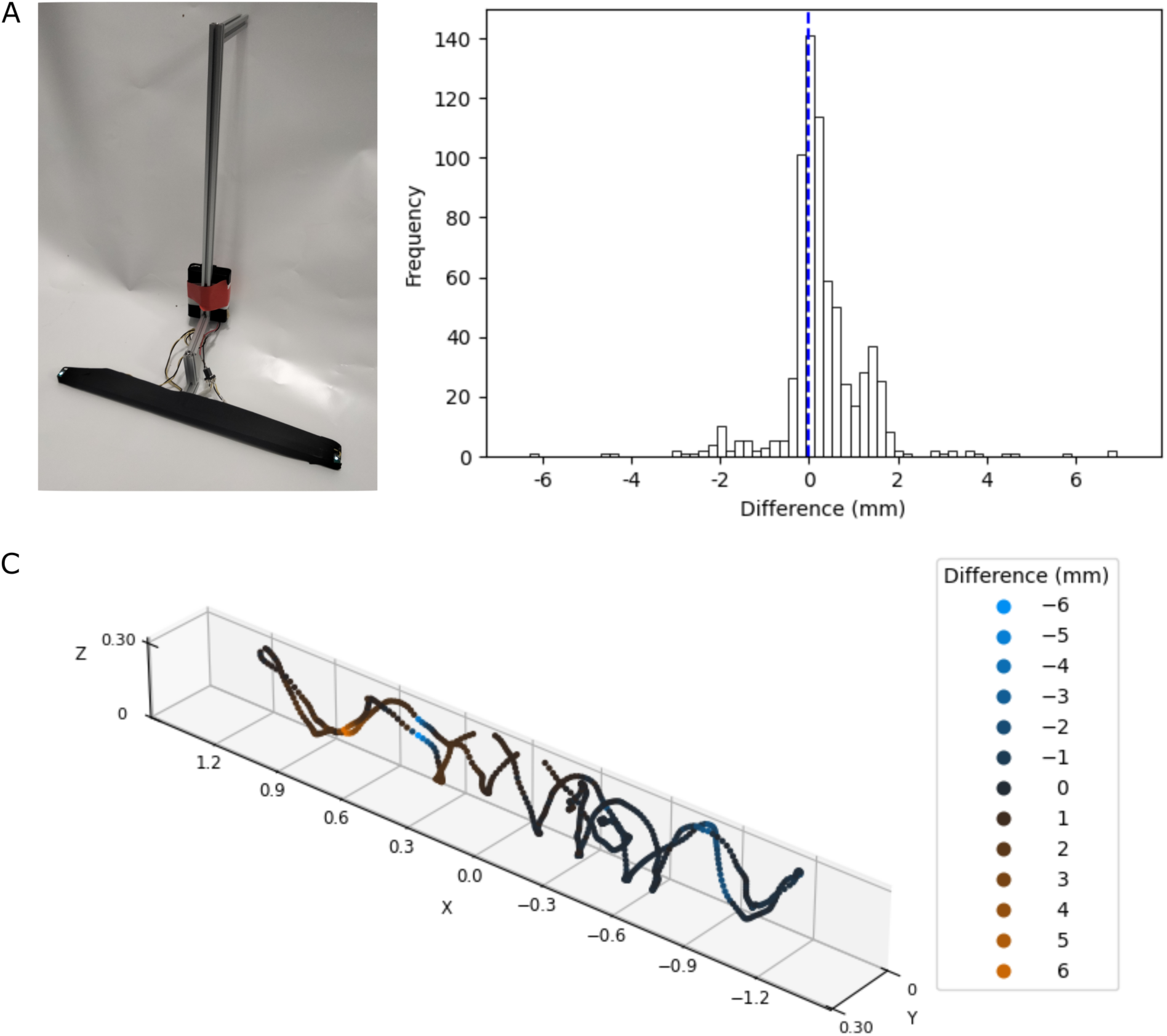
Results of the validation tests. a) The two LED wand used. b) The frequency distribution of difference between the true distance between the LEDs and Braid’s estimates within the flight arena (calculated as: *Braid’s estimate – measured value*). Here positive values indicate Braid’s estimate of the distance between LEDs is larger than the true value, negative values that Braid’s estimate is shorter. Dashed blue line indicates a difference between measurements and estimates of 0. c) a 3D representation of Braid’s distance estimates throughout the arena during validation. Axes are represented in meters and describe the flight arena’s 3D axes. Plotted are the mean position of the two LEDs according to Braid. Color of the points indicates the difference between the true distance between the LEDs and Braid’s estimates (as discussed in A). The color scale is indicated in the legend to the left of the plot: Braid estimates lower than the true value are more blue, estimates higher are more orange, darker and more black points indicate more accurate estimates of the distance between LEDs. Data used is retracked data but maintaining the tracking parameters used in config.toml (as discussed in step O2).

This protocol represents the basic setup for Braid. However, Braid is highly adaptable. Braid has been successfully applied to track insects and other relatively small targets in very complex environments [2,4,5,7], including outside environments [3,6]. The Braid system can also be adapted to track objects underwater with cameras mounted above. In our setup we describe using precision time protocol to synchronize cameras, frame acquisition can also be triggered using digital pulse to camera trigger ports using an Arduino based trigger box device (code and instructions available at https://github.com/strawlab/triggerbox). In such cases the PC communicates with the Arduino to build a live model of the clocks on both systems and allows very precise (sub-millisecond) determination of the timing of trigger pulses received by the cameras allowing precise timing information to be taken from the acquired image frames. This also allows use of USB connected cameras. Strand-Braid can run on any camera that has either the Basler pylon API (as set up here) or Vimba API (https://www.alliedvision.com/en/products/vimba-sdk/). Strand Camera object detection can be altered to fit the requirements of a given application. This includes changing the polarity of object detection to detect darker or brighter objects than the background model or, as used here, to detect absolute difference from the background (see general notes). While real time tracking is a strength of Braid, suitably synchronized video recordings can be processed in the same way offline (command ‘braid-process-video’). This can also allow re-processing of recorded video with different object detection settings (see braid’s full documentation on ‘braid-process-video’). Further documentation on these additional features is available at https://strawlab.github.io/strand-braid/.

## General notes and troubleshooting

### General notes

1. *When to recalibrate*

As noted in step G of the protocol calibration is often a more iterative process than linear protocols imply. There are several steps where changing aspects of the system may be taken that may necessitate recalibration, (i.e. repeating a calibration step after the changes have been made).

A new intrinsic calibration for a camera (step H) is required if: you change any aspect of the camera lens, such as focus, zoom, filter or the lens itself. Altering lens aperture may affect distortion of the image, this will depend on the lens but if so, such a change requires a recalibration. Note that changing the position of the camera alone does not require a new intrinsic calibration, if none of the previous camera settings are altered.

A new extrinsic calibration (step I) is required if: you have had to perform a new intrinsic calibration of any camera in the system, or you have repositioned the cameras in any way (rotation, angle, translation - xyz, position).

If you move an AprilTag to a new position and none of the above are changed in the process, you do not need to perform a new calibration, as the AprilTags movement alone does not affect the validity of the current calibration. However, you should ensure the tag’s position is adjusted in the <3DCOORDS> for when you next calibrate. It is also best practice to repeat extrinsic calibrations when AprilTags are moved to ensure the current arrangement yields a suitable calibration.

*Note: It is good practice to frequently re-calibrate the system if it is used persistently over long periods of time. Particularly as camera positions may change imperceptibly over time due to heat expansion/contraction of metal supports, the weight of the cameras on the camera mount joints or accidental movements. Typically, we perform a new extrinsic calibration (as positions of camera are more likely to be altered) every few days if the system is in use*.

*2. Changing camera frame rate in Braid*

In PTP synchronization mode camera frame rate is controlled by the ‘periodic_signal_period_usec’ parameter within the Braid configuration toml (on line 23 of the config.toml file provided). This is the interval at which cameras are triggered. So, if set at 25000 *μ*sec this will result in cameras synchronizing at 40fps. Increasing or decreasing this number will result in corresponding changes in the frame rate cameras synchronize to. This can be edited in the same manner as described in step N2.

*Warning: It is important to note if the period is set lower than the camera exposure time (controlled by the basler .pfs file), the PTP network may still be able to send trigger signals at this rate, but as cameras will not be able to collect frames with every PTP signal (as the exposure time means it will take longer than the signal period to collect a frame) this may result in the resultant frame rate differing from expectation and cameras desynchronizing*.

*3. Is Rerun a dependency of Braid?*

Braid will function without Rerun, although helpful it is not required. If one does not wish to use Rerun, omit steps corresponding to Rerun specifically (B5, I6, L3, M2) and remove the input ‘--rerun-save <rerun_save.rrd>’ from step I4.

4. *Get help with Braid and Strand Camera commands*

All Braid and strand camera commands can provide guidance on inputs. Normally if inputs are incorrect a short help message will appear in the output of the command terminal. If a full help message is required (including listing of all input options) run the command with ‘--help’ as an input, e.g.: `braid-run --help .`

5. *How to uninstall Braid*

Should you need to uninstall Braid, this can be done by running the following command in the Linux terminal:

sudo apt remove strand-braid

when doing so you may have to also manually reset the Linux package dpkg. If so, an error message will prompt you to

manually run

sudo dpkg --configure -a

before repeating the `apt remove` command. Note that uninstalling Strand Braid does <u>not</u> remove files collected by Braid, such as those saved in BRAID-DATA.

### Troubleshooting

*Problem 1: pylon Viewer (and pylon IP configurator) will not launch after installation and the associated steps have been properly performed*.

Possible cause: There is a problem with how pylon API lists its Linux package dependencies. This means some software packages that pylon API requires to run are not always installed (usually the ‘libxcb-xinput0’ dependency of the package ‘libqxcb.so’ is missing – but detailed below are instructions on identifying missing dependencies).

It is worth noting the packages whose omission prevents pylon Viewer from launching are needed for pylon Viewer itself, not strand-braid. Thus, if the installation is otherwise conducted as described, the components of the pylon API needed to run strand-braid should be installed and function normally. However, difficulties may arise from not being able to access pylon Viewer and the IP configurator.

To confirm that this dependency issue is the cause try running the command:

/opt/pylon/bin/pylonviewer

This command launches pylon Viewer via the command terminal. If this results in the following output this dependency error is probably the case:

qt.qpa.plugin: Could not load the Qt platform plugin “xcb” in “” even though it was found.

This application failed to start because no Qt platform plugin could be initialized. Reinstalling the application may fix this problem.

Available platform plugins are: eglfs, linuxfb, minimal, minimalegl, offscreen, vnc, wayland-egl, wayland, wayland-xcomposite-egl, wayland-xcomposite-glx, xcb.

Aborted (core dumped)

Solution: Identify the packages that are missing and install them. The following steps will allow you to do this.

1. Attempt to launch pylon Viewer with the QT debug information:
a. Turn on the QT debug output using the following command

export QT_DEBUG_PLUGINS=1

b. run pylon Viewer in the command terminal using the following:

/opt/pylon/bin/pylonviewer

c. The output of this will look like figure 10, where various plugins and whether success has been achieved getting the ‘keys’ from these will be listed (a successful plugin, ‘eglf’, is shown in figure 10a). Usually, the plugin that has failed to load is ‘xcb’ (the failed plugin, ‘xcb’, is shown in figure 10b). The output here will identify a path to a ‘.so’ (shared object) file that failed and causing the aborted launch of pylon Viewer. This can be identified on the output, in figure 10b. Usually it is the .so object at this path:

**Figure 10:**
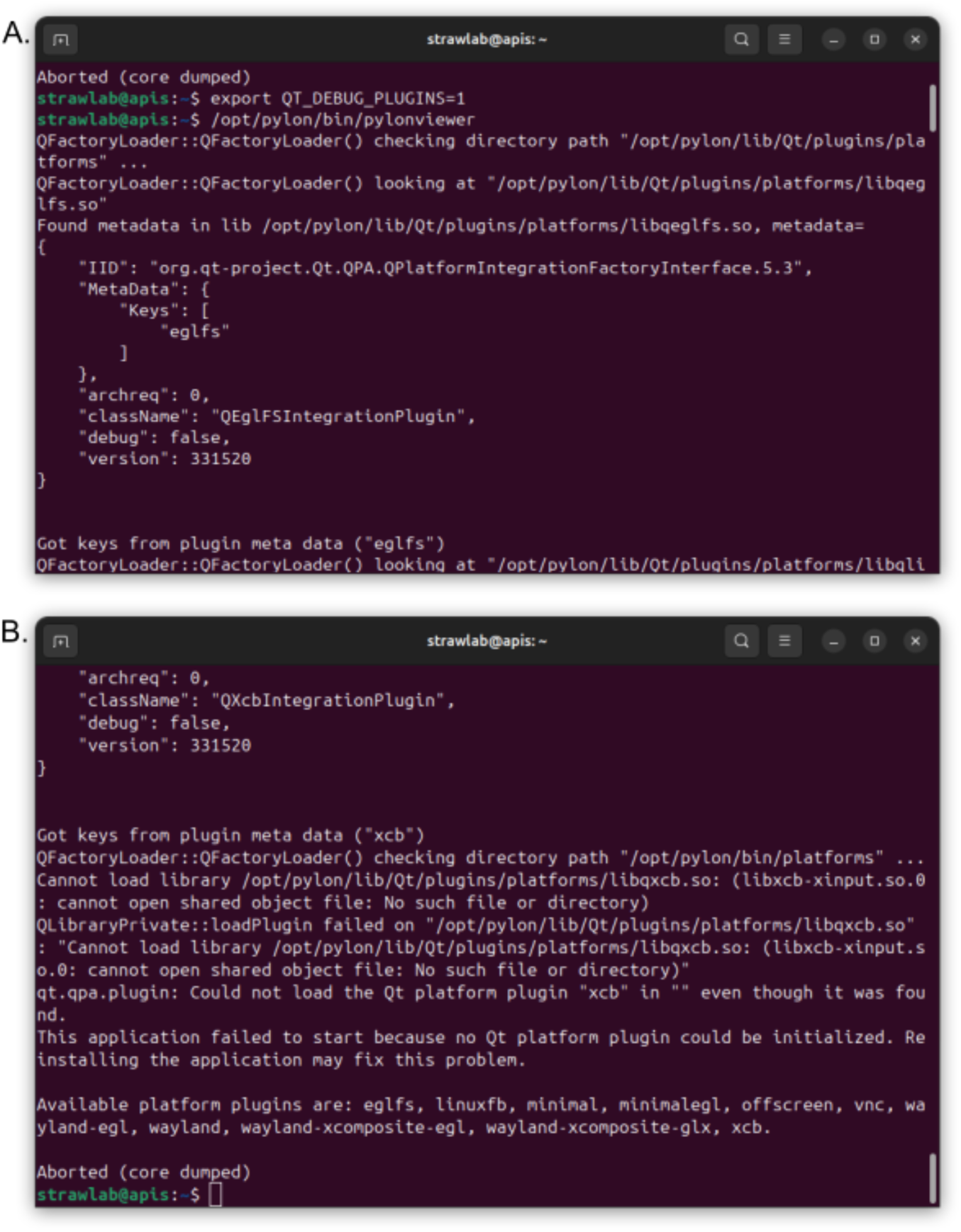
a subset of the Qt debug output when pylon Viewer is launched, many plugins will be listed here but each should have outputs will be similar to either panel. A. a successful plugin, ‘eglf’, output. B. a failed plugin output, shown for ‘xcb’ the plugin whose dependencies usually needs to be installed manually. Note the path to the .so file related to this plugin.

/opt/pylon/lib/Qt/plugins/platforms/libqxcb.so

2. Identify the dependencies of this ‘.so’ file. By running:

ldd <SO_FILEPATH>

where <SO_FILEPATH> is the path to the failed ‘.so’ file identified in above. If the failed ‘.so’ file were libqxcb.so:

ldd /opt/pylon/lib/Qt/plugins/platforms/libqxcb.so

The output of this command will list several packages followed by ‘=>’ indicators, and file paths to their ‘.so’ files. These are the dependencies of libqxcb.so. Identify the packages which are missing, these will not have a file path and will be listed as ‘not found’ following the ‘=>’

3. Install those packages using the following command:

sudo apt-get install <PACKAGE_NAME>

where <PACKAGE_NAME> is the name of missing packages.

### Note: package names for apt-get do not always exactly match the package name once installed. Use apt-cache search

*<SEARCH term>* (where *<SEARCH term>* might be the package name, or part of it) *to identify the package name for apt-get*.

Usually the following is required:

sudo apt-get install libxcb-xinput0

Once this step is complete, the output of the command given in problem 1 step 2 should not list any dependencies as ‘not found’, if this is the case pylon Viewer should now launch. You may be required to restart the PC first. If problems persist consult Basler customer support: https://www.baslerweb.com/en/learning-support/.

*Problem 2: The following Error appears onscreen: “Error: frame processing too slow, Processing of image frames is taking too long. Reduce the computational cost of image processing.”*

Possible cause: The current task Braid is carrying out is computationally demanding such that it is not successfully performing the task on every frame received. This normally happens during checkerboard calibration (step H) or AprilTag detection (Step I).

Solution 1: During checkerboard calibration this error can be safely dismissed. Doing so will simply result in checkerboards being detected at a much lower rate than that of the camera’s set frame rate. Just ensure the checkerboard counter at *Strand Camera→ Checkerboard Calibration → Number of checkerboards collected* ticks up once the checkerboard is in the correct positions (as noted in step H3).

Solution 2: Collection of checkerboard and AprilTag detections can be conducted in individual instances of Strand Camera launched directly (as in step E) as well as via instances accessed via Braid (step F4). Performing these steps via individual instances of Strand Camera launched directly one at a time can be less demanding.

Solution 3: Reduce the frame rates of cameras for the current task see General notes 1. For AprilTag detection and checkerboard calibration it is fine to run Braid at 10FPS

*Problem 3: Cameras do not synchronize and Braid’s activity log gives the warning that “launch time precedes device timestamp. Is time running backwards?” continuously*.

Possible cause: ptpd is improperly configured or not started. This warning will occur either when ptpd is not available and Braid attempts to synchronize cameras with the less accurate network time protocol (NTP) which will fail to meet the set tolerance for synchronization, or when the PC clock is not set as the ‘Master’ clock for the PTP network, this can lead to one of the camera’s clocks being used as the ‘Master’ and is ahead of the PC’s clock (i.e. the clock used by Braid).

Solution: Ensure ptpd is functioning properly. Repeat step B3.

*Problem 4: GigE cameras were previously found by Strand-Braid. These cameras have not been disconnected, but now Strand-Braid and pylon Viewer states ‘Cameras not found’*.

Possible cause 1: Gigabit network connection has been deactivated, this can happen when the switch but not the PC loses power, or when newer network connections are made such as Wi-Fi or a regular Ethernet connection to the PC. This can cause the PC to disable ‘older’ connections in favor of new.

Solution 1: Ensure the Gigabit and ethernet cable connections are in order and that the switch is powered. Ensure the PC is connected to the switch and external internet as in step A, and Wi-Fi is not connected. Ensure the Gigabit network connection is activated by repeating step A4.

Possible cause 2: A network error has occurred, and the cameras have not connected. Open the *pylon IP configurator*

application. Under the column *Status* the cameras will read ‘unreachable’ if this is the case.

Solution 2: Refresh the Gigabit and basler IP settings

a. Go to *pylon Viewer* select *Tools→pylon GigE configurator* and select the *Configure* button in the window that appears. You will be asked for you password, enter it and allow the *pylon GigE configurator* to finish (this should only take a second).
b. Open the *pylon IP configurator* if cameras do not display ‘OK’ as in figure 1, and still display ‘unreachable’ refresh each camera’s IP settings.
i. Check the IP configuration and IP Address of each camera.
ii. If IP configuration is incorrect for your network, change the settings using the selections on the bottom left of the window and click save.

*Hint: consult your network manager for clarification of the network’s IP protocols*.

iii. If IP configuration is correct but the IP Address incorrect, refresh the IP configuration of the camera by selecting a protocol configuration in the bottom left of the window, then click back to the correct IP configuration mode and then save (that is save the configuration without having made a change). Upon the save the camera should retry the connection and be detected.

*Problem 5: Launching strand camera or Braid results in a ‘BackendError(Opening camera)’ the output of which states, in addition to other things, that ‘The device is controlled by another application’*

Possible cause: the camera is currently open in another application, most likely another strand-Braid instance started previously is still running or pylon Viewer is running with a camera accessed. The pylon API allows only one program to operate (that is ‘open’) a camera at a time. This is why it is good practice to close such applications that access the cameras once finished with them.

Solution: identify and close the application already using the camera and start strand-braid again, the most likely candidates are pylon Viewer or another Strand-Braid instance that has not been properly stopped.

## Supplementary information

A single Supplementary materials file is provided that contains Supplementary materials 1: PC specifications, Supplementary materials 2: Additional Equipment Diagrams and Supplementary materials 3: Additional bee tracking figures.

## Videos

**Video 1:** A screen recording demonstrating launching Strand Camera. Keys and mouse clicks are indicated by yellow text in the bottom left of the screen. First Strand Camera is launched (at 00:05) without specifying the camera name (as in D2). If launched in this way a Strand Camera instance is launched using the first camera detected (note that if following the protocol to step D2, this first detected camera will be the only camera connected to the PC). When strand Camera is launched a GUI window is opened in the browser (at 00:10). Looking at the Strand Camera GUI (pointer indicator at 00:12), or in the activity log (pointer indicator at 00.19), we can identify the system camera name. We can then launch Strand Camera on this camera specifically as we do the second time strand camera is launched (at 00:50, as in step E3). Note that if the GUI is closed but the terminal command is still running (as performed at 00:57), Strand Camera does not stop running and the GUI for the strand camera instance can be accessed again by returning to the same http address (which can be found in the terminal log, as performed at 01:00). Strand Camera instances can be stopped in the terminal (e.g. closed or interrupted with *Ctrl+c* as at 00:39) or by clicking the ‘Quit Strand Camera‘ button (as done at 01:11). When the terminal command for an instance of Strand Camera is stopped (e.g. closed or interrupted with *Ctrl+c* as at 00:39) this stops the current Strand Camera instance, subsequently leading to the GUI losing connection to Strand Camera.

**Video 2:** A screen recording of launching Braid. Keys and mouse clicks are indicated by yellow icons in the bottom left of the screen. Braid is launched from the Linux terminal (00:07). As this is the first time the system has been launched since the cameras were turned on it takes some time to synchronize so, we wait until confirmation occurs in the log that cameras are synchronized (00:16), we then scroll up to the URL for the GUI (accessed at 00.27). Upon accessing the Braid GUI, by clicking the individual camera names we can access the individual Strand Camera instances making up this Braid instance (00:31-01.20). Performing *Ctrl+c* in the terminal running Braid will stop this instance (01:40), interrupting the connected GUI (as Braid is not running). Braid can also be closed by clicking the ‘Quit Braid’ button (as done at 2:21). Note that due to the port being specified in the configuration toml (input *‘*http_api_server_addr*’*) if Braid is relaunched with the same toml (as in 01:45) the GUI will be accessible at the same address as before.

*Note: video 2 is launched with a config file containing a calibration. If Braid is launched without a calibration file the ‘calibration:’ indicator in the Status tab will in the Braid GUI will read ‘NO CALIBRATION’*.

**Video 3:** A screen recording of performing checkerboard image collection procedure and intrinsic calibration for a single camera. Keys and mouse clicks are indicated by yellow icons in the bottom left of the screen. To facilitate checkerboard collection view is split between two tabs one showing the *Live view* and the other the *Checkerboard Calibration* tabs (00:05). This screen view is then made visible to the experimenter while checkerboards are collected. Checkerboard size inputs are then confirmed (00:11, step H2b). Here the defaults are correct. Then checkerboard detection is enabled (00:14, step H2c), as is ‘save debug information’ (00:30). The checkerboard is shown to the camera at each position listed in table 1 (00:50-02:23, step H3). At each position the checkerboard is held until detection is confirmed on the counter at *Checkerboard Calibration → Number of checkerboards collected* before a new position is assumed. In this example 17 checkerboard images in total are collected (02:49 this counter indicated). Once checkerboards are collected, detection and saving debug information are turned off (02:53). The calibration is then produced (3:00, step H4a) and the output calibration is navigated to (03:05, step H5a). Observe that two calibrations are produced (the calibration itself and a timestamped copy, Basler-81011970.yaml, indicated by the pointer and opened at 03.21, and Basler-81011970_140111.yaml). The calibration’s mean reprojection distance is checked, here it has a suitably low value of 0.53 pixels (see step H5b). Similar videos for all cameras are available in [77].

**Video 4:** The Rerun multicamera and 3D view while tracking a single bee (the bee with the time identifier ‘1427’, see the datasets included with this publication). The appearance of Rerun when running live or when a rrd file produced by braid is viewed is similar. Rerun presents bee position via a dots placed on the camera views and 3D visualization. Within the 3D visualization the arena bounds are indicated by a simple 3D model (provided in [77] that can be imported into Rerun viewer). A static version of this bee’s track is provided in figure 6.

## Acknowledgments

This work was funded by the Volkswagen Foundation Momentum Program (98 692 to A.D.S.).

We thank M. Siegel and the mechanical workshop of the Institute of Biology I. N. Brehm, and T. Albonetti provided bee keeping services.

## Ethical considerations

No ethical permissions were required for the experiments involving bees, but the experiments were conducted according to ASAB/ABS guidelines.

