## Supplementary materials for "Low-latency multicamera 3D tracking of insects with Braid"

### Low latency multicamera 3D tracking of insects with Braid, Supplementary materials:

#### Supplementary materials 1: PC specifications:

Below are the critical components related to Braid performance of the Linux PC used in the protocol. Other hardware (tower case, USB keyboard, mouse, HDMI screen) is also required but their specifications will not impact Braid functionality.

**Motherboard:** ASRock, Taipei, Taiwan **model:** X570S PG Riptide

**CPU:** AMD, Santa Clara CA, USA **model:** Ryzen 7 5800X 8-core

**Graphics card:** NVIDIA, Santa Clara CA, USA **model:** GP106GL [Quadro P2000],

with Graphics driver: nvidia v: 550.163.01

**RAM/Memory:** Kingston Technology, Fountain Valley CA, USA **model:** 9905734-415.A00G type: DDR4 size: 16 GiB speed: 3200 MT/s

*Note: Two of the listed RAM cards were used in the PC*

Appropriate CPU and PC tower fans and heat sinks are also required as per your lab conditions. Additionally, sufficient hard drive data storage is required if video is also to be saved, the PC used in the protocol had a 2TB hard drive. This is far more than that required for the data collected in the protocol (detailed in main text).

#### Supplementary materials 2: Additional Equipment Diagrams

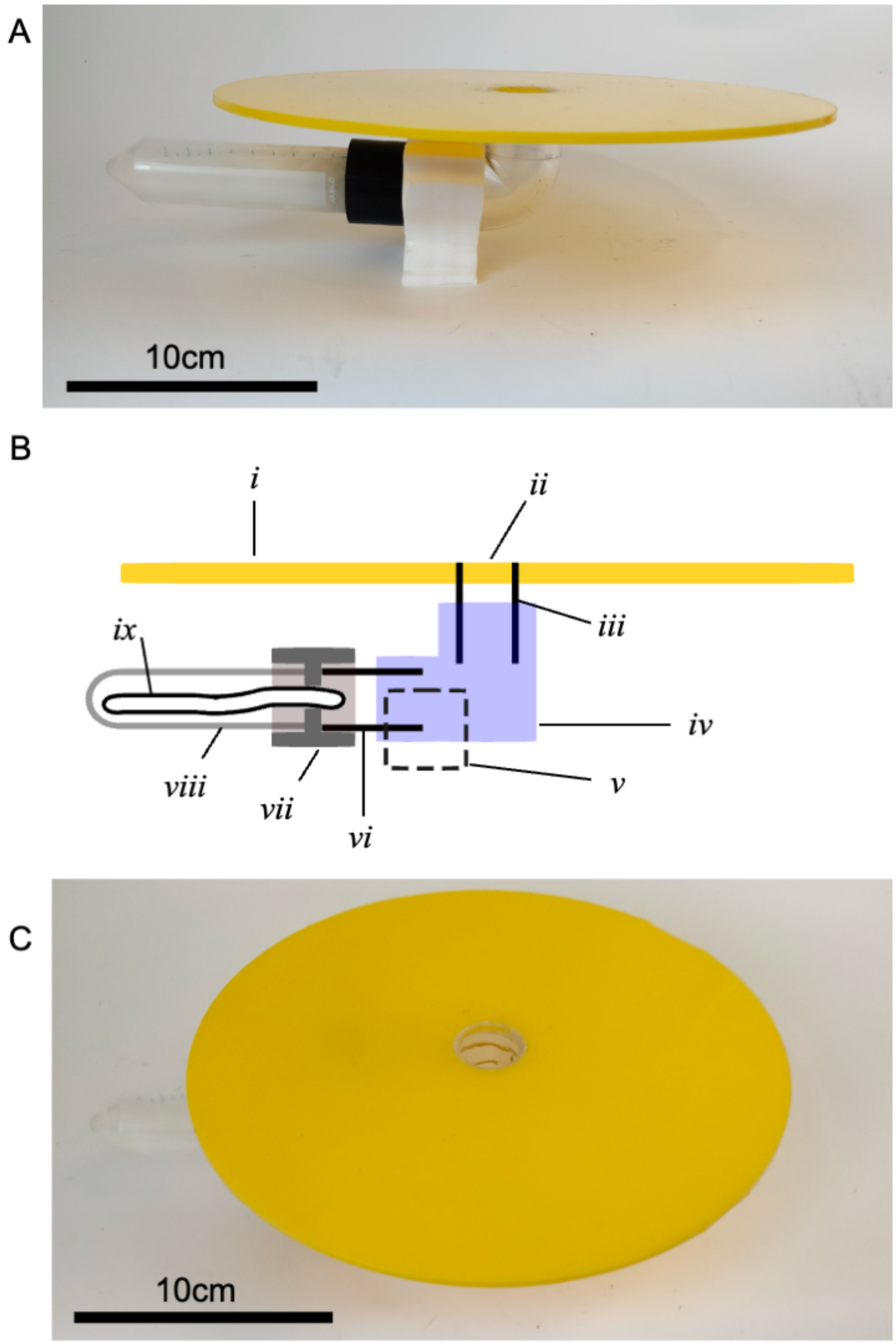


**Figure S1:** the 'type A' artificial flowers. A) photographic image of the artificial flower in profile. B) diagrammatic representation of image A, note that this diagram is not to scale. Italic numerals indicate components and features. i, The yellow acrylic disc. ii, The entrance hole to the flower. iii, a 5cm section of 32mm diameter pipe. iv, a 90-degree pipe fitting for 32mm pipe. v, polystyrene stand. vi-ix, together indicate the falcon tube collar and wick design feeding reservoir, consisting of: a 5cm section of 32mm diameter pipe (vi) that is inserted into a 3D printed flower collar (vii) that links the pipe to a falcon tube filled with sucrose solution (viii), through which a wick (ix) is inserted. C) the artificial flower as it appears from above. Note scale bars in photographs are approximate due to image perspective view.

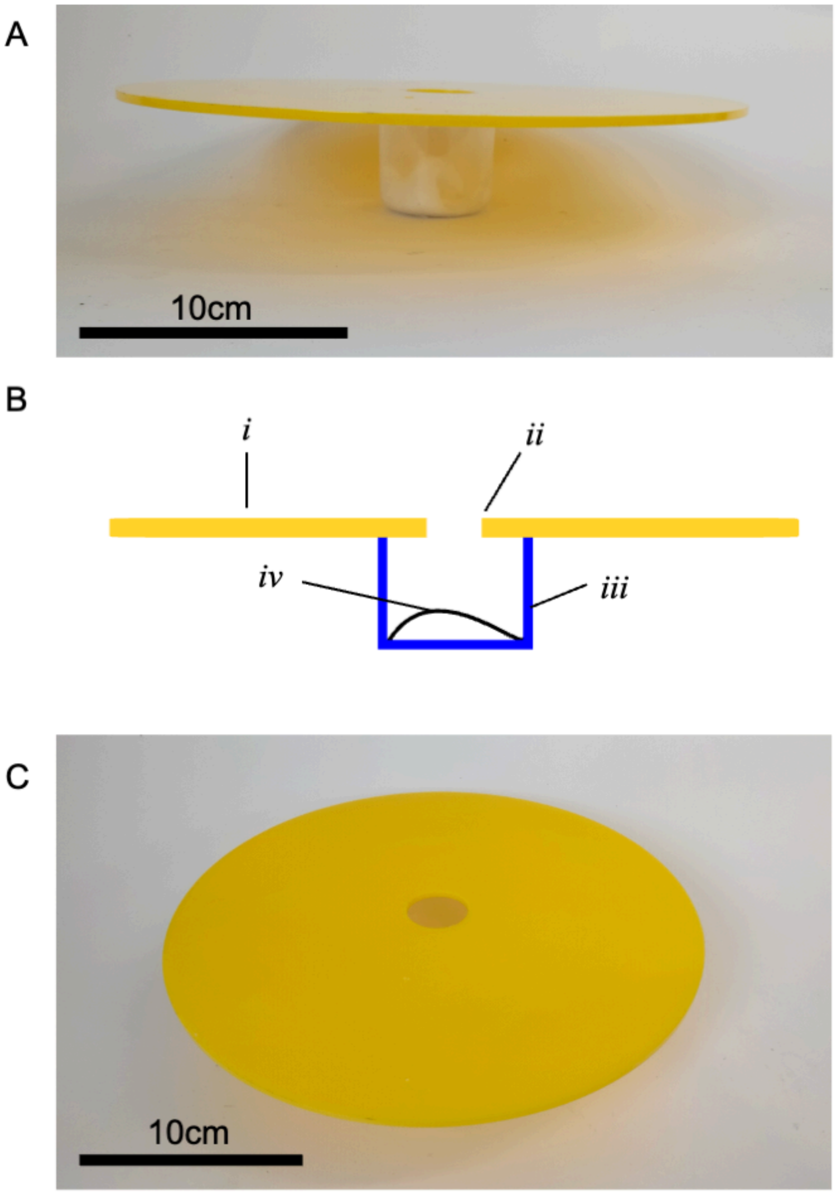


**Figure S2:** the ‘type B’ artificial flowers. A) photographic image of the artificial flower in profile. B) diagrammatic representation of image A, note that this diagram is not to scale. Italic numerals indicate components and features. i, The yellow acrylic disc. ii, The entrance hole to the flower. iii and iv together indicate the plastic jar design feeding reservoir, consisting of: a plastic jar (iii) and a cotton wool wick soaked in sucrose (iv). C) the artificial flower as it appears from above. Note scale bars in photographs are approximate due to image perspective view.

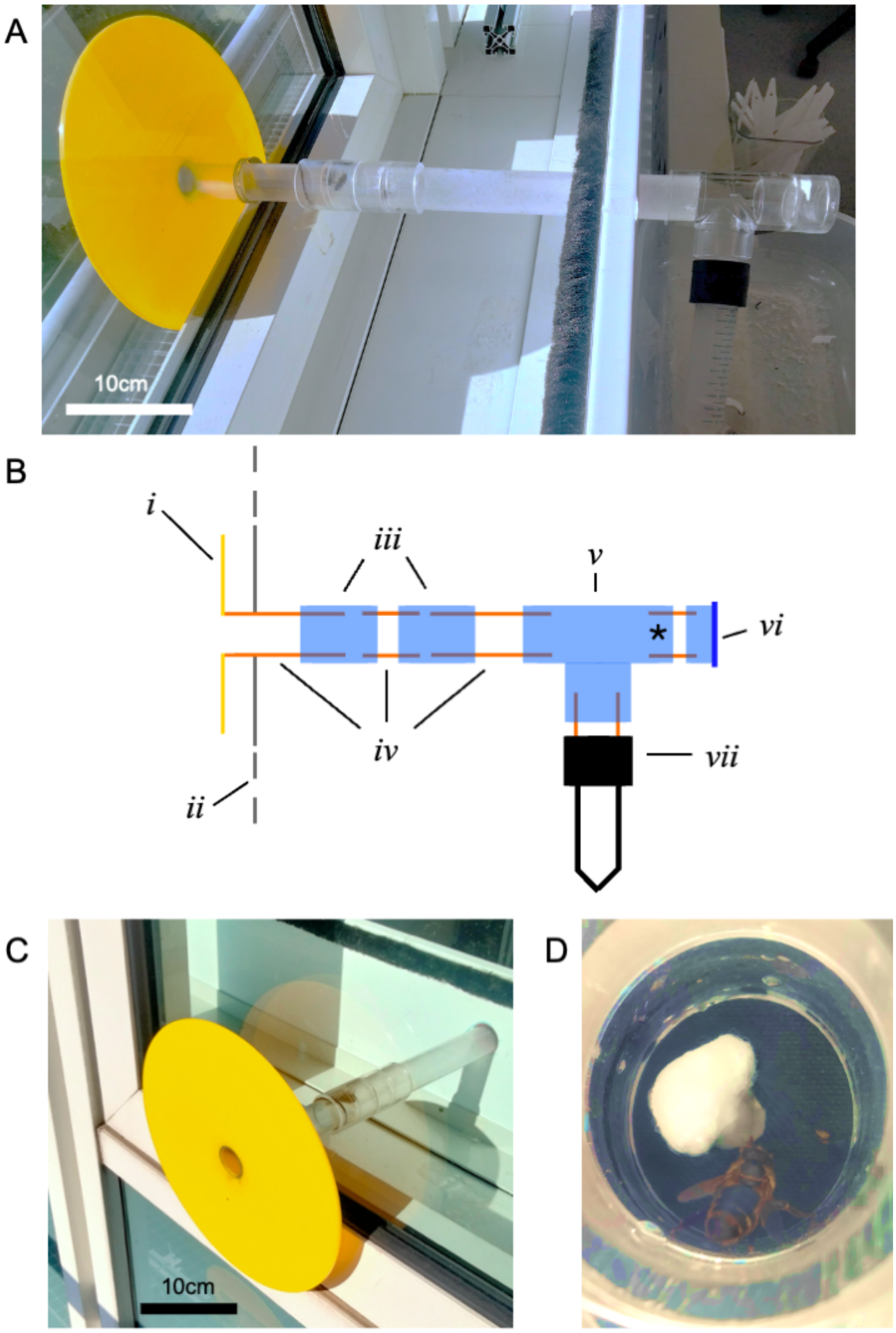


**Figure S3:** The window-access when access to the flight arena is blocked. A) photographic image. B) diagrammatic representation of image A, note that this diagram is not to scale. Italic numerals indicate components: i, an artificial flower disc. ii, the windowpane. iii, horizontal pipe connector fitting for 32mm tubing. iv, 32mm diameter piping (lengths left to right: 10cm, 5cm, 30cm). v, T-junction pipe connector for 32mm tube. vi, a 5cm length of 32mm pipe that leads to a closed cap. vii, Falcon tube feeding reservoir. Note between items iii. and v., there is a hole drilled into the windowsill through which piping passes for support. This is not shown in the diagram. The point indicated by '*' indicates where the arena access tube connects to the window access. In A and B this connection is blocked by a cap (vi), as at the start of the protocol. This point is also indicated in figure S4B by the same symbol. C) exterior view of the Lab-window access. D) A honey bee feeding from the wick in the falcon tube collar of the Lab-window access. The complete width of D is approximately 3cm. Note that photographic images have been sharpened and color adjusted to clarify the view of the key features. Note scale bars in photographs are approximate due to image perspective view.

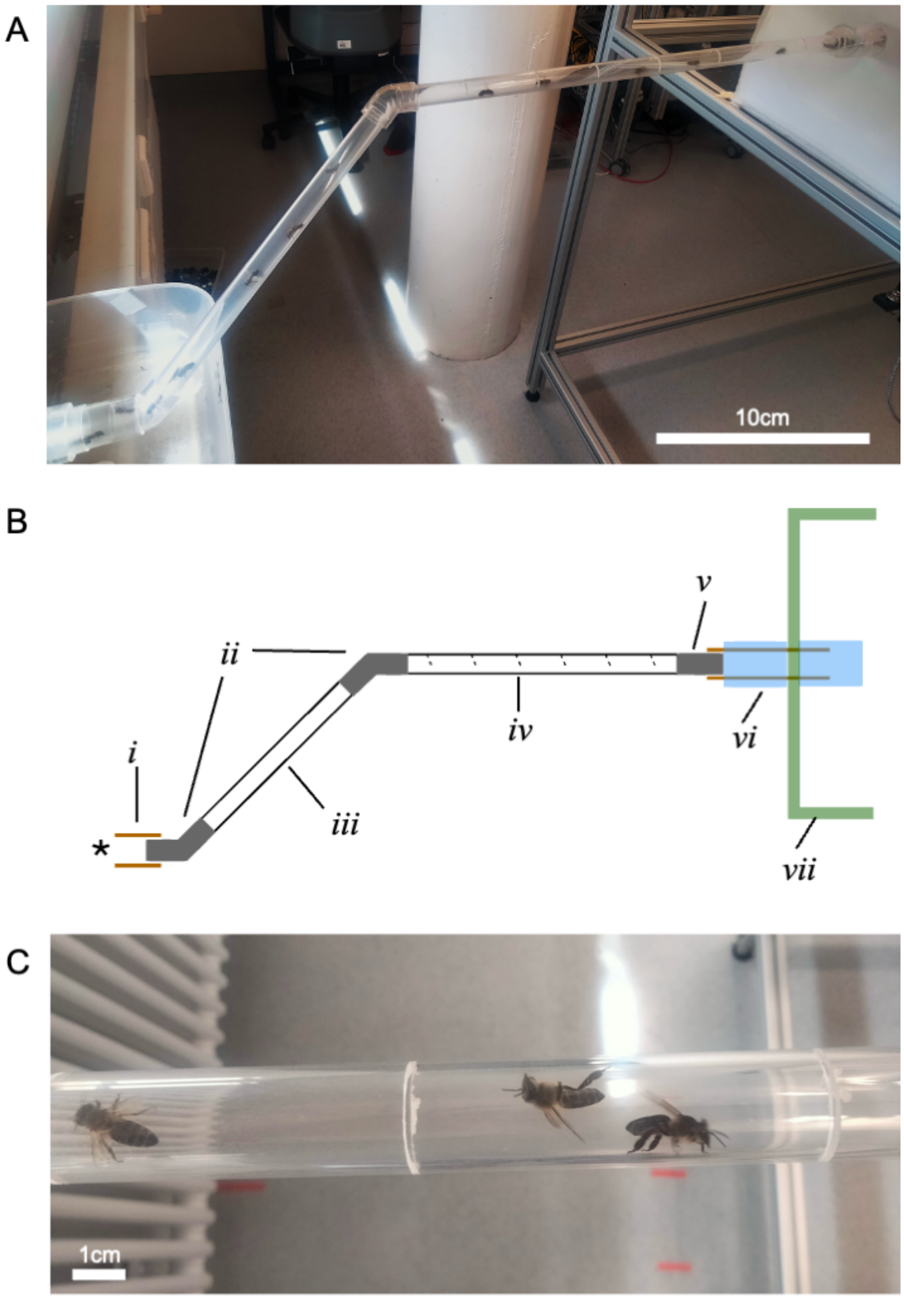


**Figure S4:** The arena access tube. A) a photographic image (image sharpened and contrast increased for clarity), B) a diagrammatic image of the access tube. Italic numerals indicate components. *i,* a 5 cm length section of 32mm diameter tube, inserted into the T junction of window access (that makes up vi in figure S3). Point indicated by '*' indicates where the arena access tube connects to the Lab-Window access, at the point also indicated in figure S3B by the same symbol. *ii,* 45-degree pipe fittings for 20mm diameter tube. *iii,* a 30mm length of 20mm diameter tubing. *iv,* a 50mm length of 20mm diameter tubing with slits cut along its width. *v,* horizontal pipe connector fitting for 20mm tube. *vi,* a 10 mm section of 32mm diameter pipe inserted through the flight arena hole (see main text) with two horizontal pipe fittings fit over either side to lock the tube in place. *vii,* the flight arena. C) honey bees within the lab access tube. Note that photographic images have been sharpened and color adjusted to clarify the view of the key features.

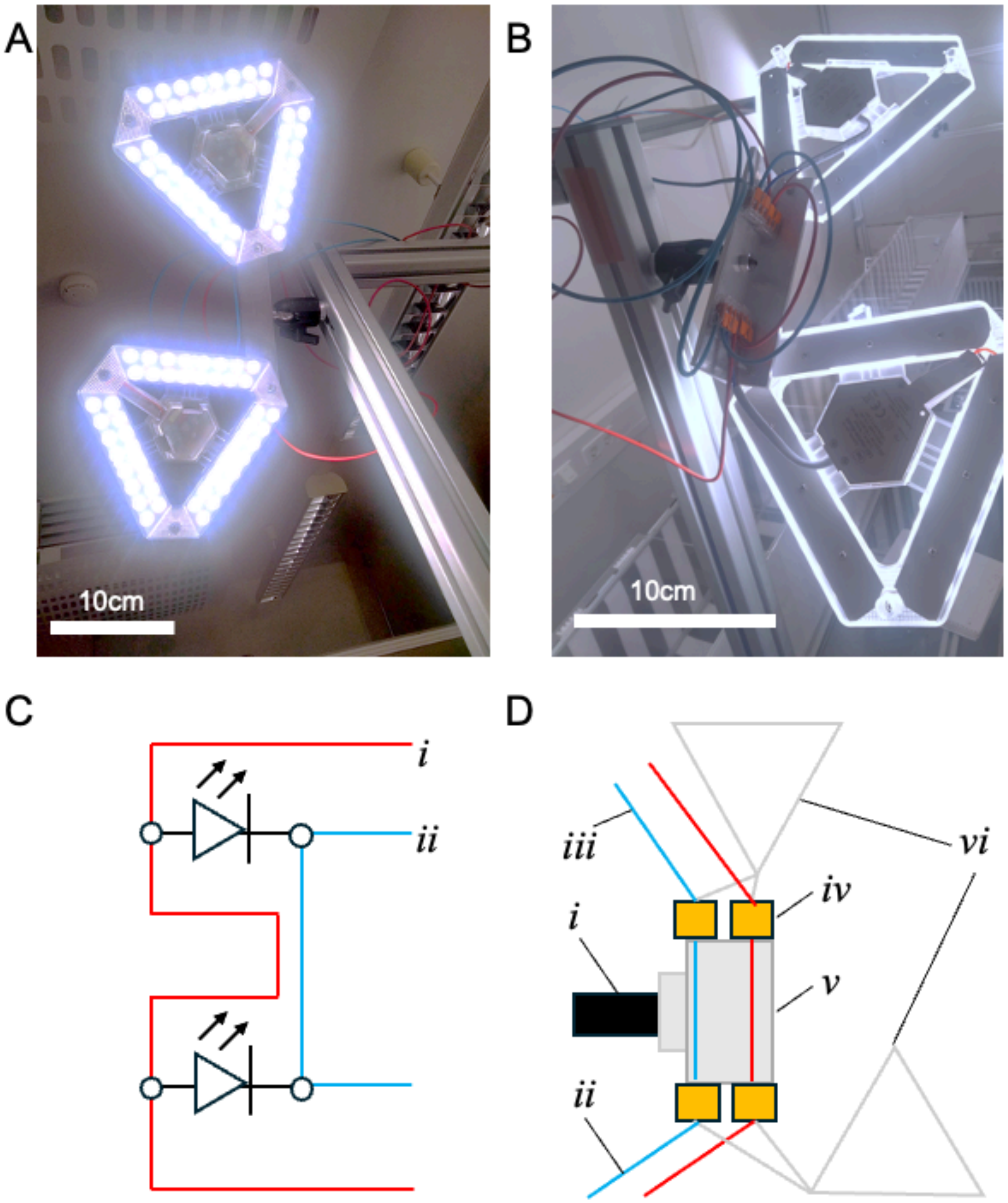


**Figure S5:** The light arrays used in the protocol. A) the array from in front. B) the array from behind. C) a circuit diagram of each array, where each LED unit is represented by a single LED, red wire the ‘positive’ wiring and blue ‘negative’ and round circles indicate the ‘Wago’ Connectors. Four such units are used. See figure S6 for detail on wiring of units together and arrangement. Note that wires terminating in *i* and *ii* are not required for the last unit, see figure S6. D) a diagrammatic representation of B. Italic numerals indicate components. *i,* the ball mount. *ii,* incoming positive wire, red, outgoing negative wire, blue. *iii,* outgoing positive wire, red, incoming negative wire, blue. *iv,* ‘Wago’ connectors. *v,* L shaped metal plate that screws into ball joint and to which connectors are stuck. Note that, positive and negative wires span the plate between connectors. *vi,* two LED light units. Note that photographic images have been sharpened and color adjusted to clarify the view of the key features.

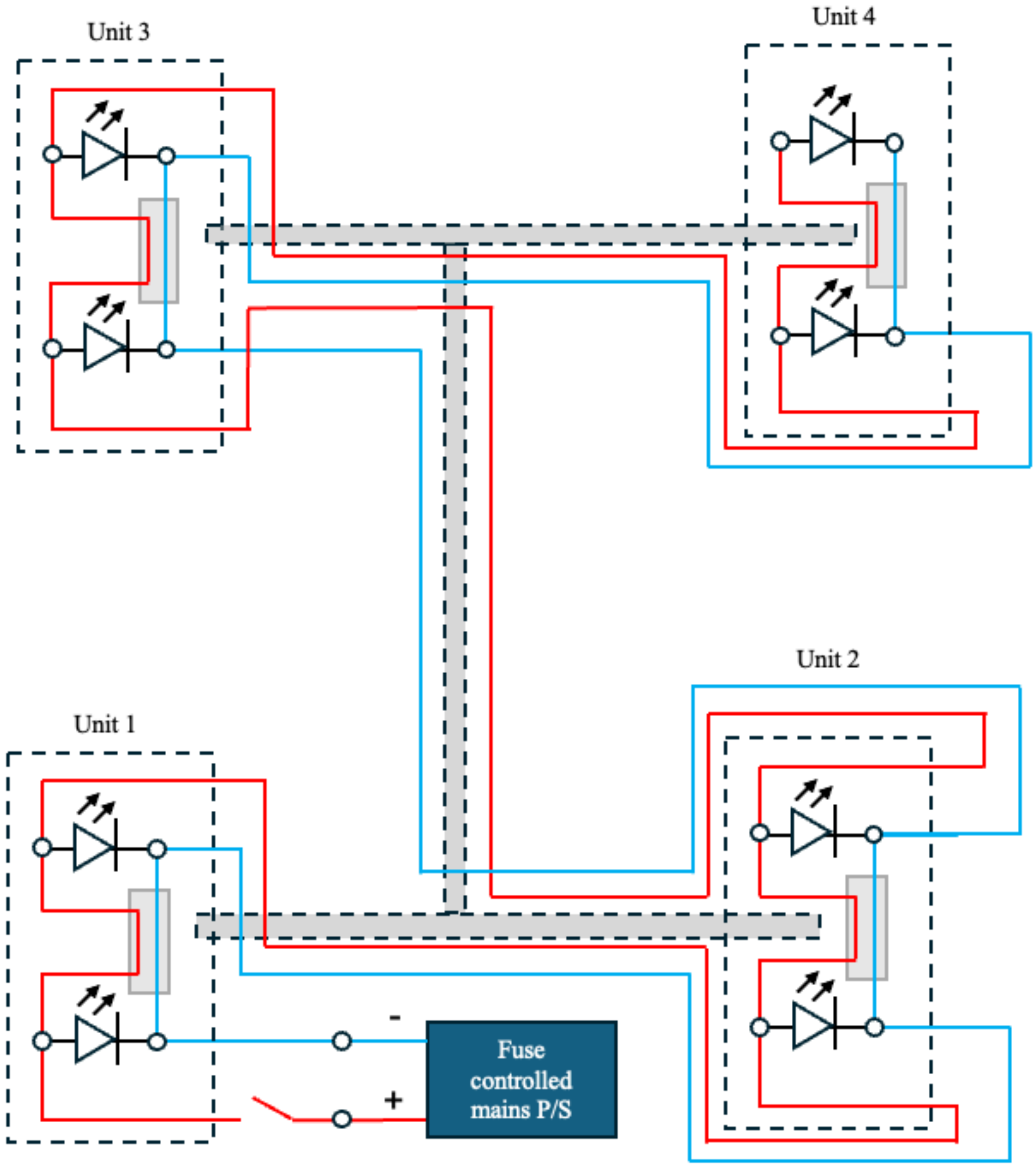


**Figure S6:** Simple circuit diagram and sketch of light array positions across the Support and camera mounting frame, where each LED unit is represented by a single LED, red wire the 'positive' wiring and blue 'negative' and round circles indicate the 'Wago' Connectors. Note that there are 4 light array units (shown in figure S5), each with 2 LED units and that Unit 4, the unit at the end of the circuit, lacks incoming negative and outgoing positive wires. Solid grey rectangles represent the mounting plate of each unit. Dashed rectangles represent the Support and camera mounting frame viewed from above. Note diagram is to demonstrate wiring of components and is not to scale.

### Supplementary materials 3: Additional bee tracking figures

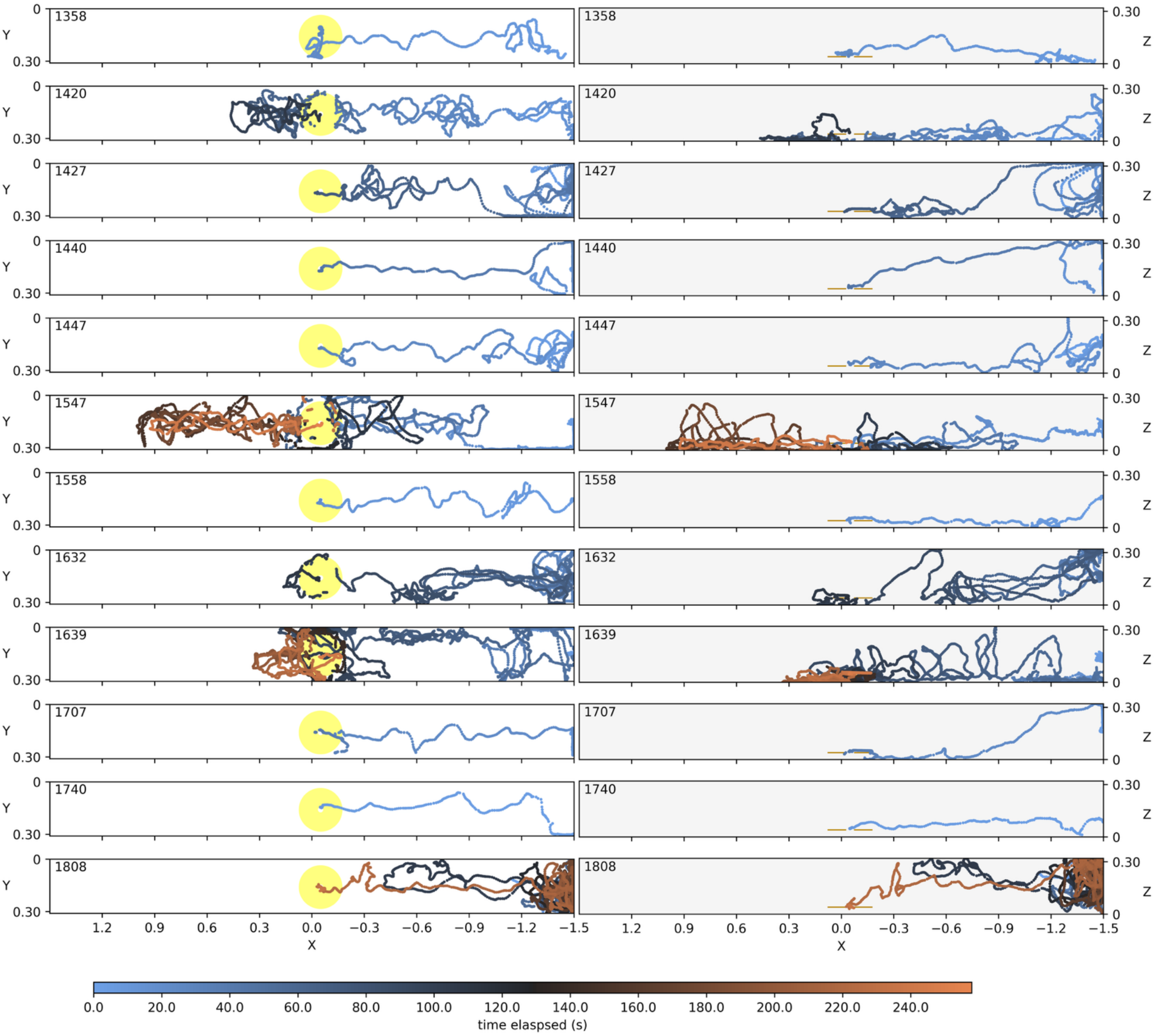


**Figure S7:** 3D tracking results of the 12 bees that successfully located the flower, retracked using the original tracking parameters given in config.toml. Each row of graphs shows the tracking for an individual bee. Individual bees are identified by a 4-digit code corresponding to the 24h time when tracking began. This ID is shown in the top left of each panel (this time can be used to identify the BRAIDZ file as all data in this step was collected on the same day). Graphs show for each bee tracking results throughout the tracking (Braid position estimate) within the x-y (left-hand side, white background panels) and x-z (right-hand side, grey background panels) axes. Axes are in meters and are as described for the tracking arena. Color of points indicates time elapsed (in seconds) since the bee entered the arena. This color scale is shared across all graphs. The artificial flower position during tracking is indicated by the yellow disc (xy graphs) or lines (xz graphs). As discussed in the protocol, note that Braid cannot track the bee once it is sufficiently inside the artificial flower or under the artificial flower's disc. Note that tracking ends when the bee enters the artificial flower. Also note how bees vary in the time they take to locate the flower.

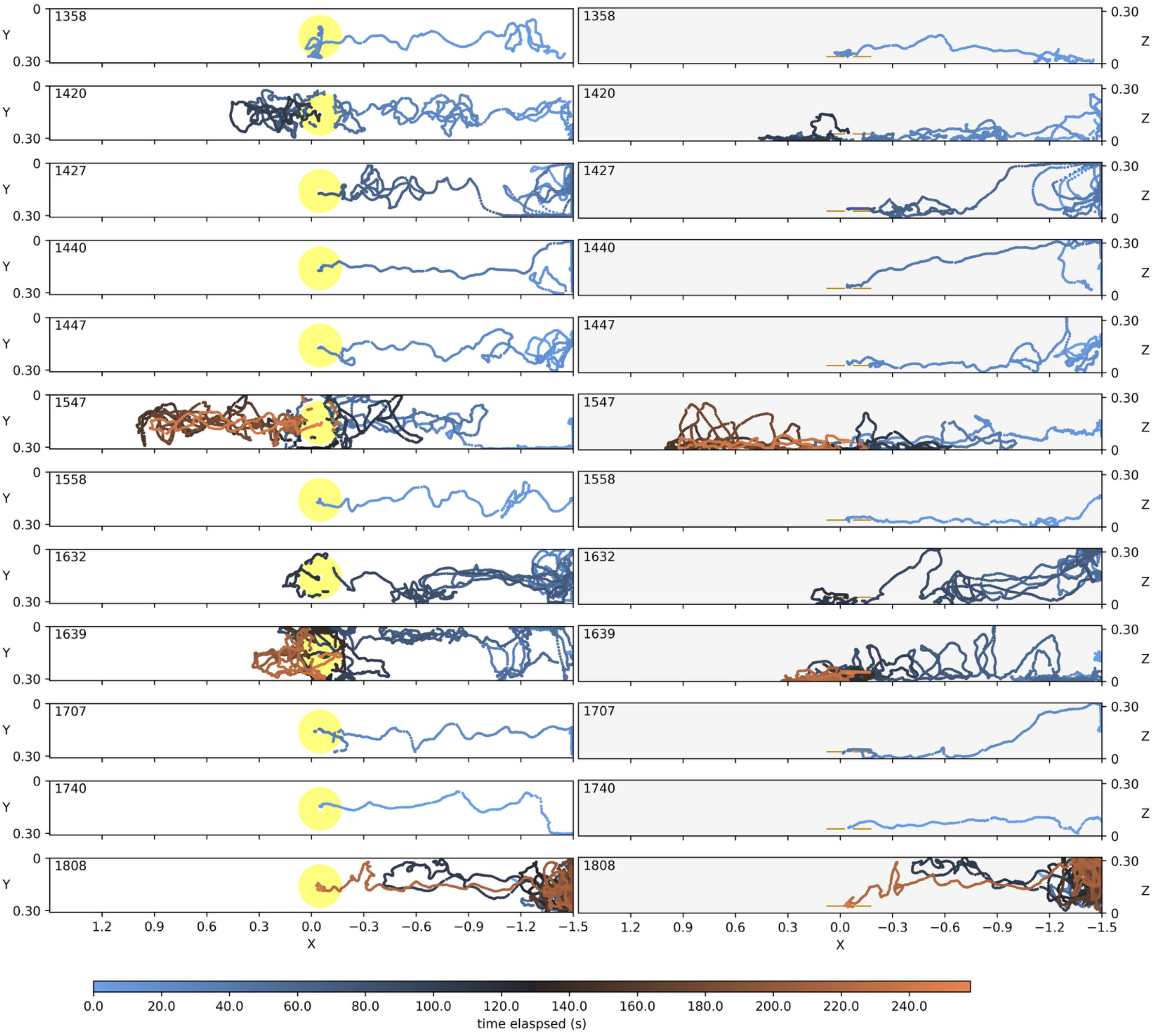


**Figure S8:** 3D tracking results of the 12 successful bees after being retracked by Braid using the alternative tracking parameters provided in retrack_paras.toml. For all aspects of the graph, see the legend of figure S7.
